# African elephants address one another with individually specific calls

**DOI:** 10.1101/2023.08.25.554872

**Authors:** Michael A. Pardo, Kurt Fristrup, David S. Lolchuragi, Joyce Poole, Petter Granli, Cynthia Moss, Iain Douglas-Hamilton, George Wittemyer

**Affiliations:** Department of Fish, Wildlife, and Conservation Biology, Colorado State University; Fort Collins, CO, USA; Department of Electronic and Computer Engineering, Colorado State University; Fort Collins, CO, USA; Save The Elephants; Nairobi, Kenya; ElephantVoices; Sandefjord, Norway; Amboseli Elephant Research Project; Nairobi, Kenya

## Abstract

Personal names are a universal feature of human language, yet few analogs exist in other species. While dolphins and parrots address conspecifics by imitating the calls of the addressee ^1,2^, human names are not imitations of the sounds typically made by the name’s owner ^3^. Labeling objects or individuals without relying on imitation of the sounds made by that object or individual is key to the expressive power of language. Thus, if non-imitative name analogs were found in other species, this could have important implications for our understanding of language evolution. Here, we show that wild African elephants address one another with individually specific calls without any evidence of imitating the receiver’s vocalizations. A random forest model correctly predicted receiver identity from call structure better than expected by chance, regardless of whether the calls were more or less similar to the receiver’s calls than typical for that caller. Moreover, elephants differentially responded to playbacks of calls originally addressed to them relative to calls addressed to a different individual, indicating that they can determine from a call’s structure if it was addressed to them. Our findings offer the first evidence for a non-human species individually addressing conspecifics without imitating the receiver.

## MAIN TEXT

One of the hallmarks of spoken human language is the use of vocal labels, in which a learned sound refers to an object or individual (the “referent”) ^4^. Many species produce functionally referential calls for food and predators ^5,6^, but the production of these calls is typically innate ^7^. Learned vocal labels allow for much more flexible communication than innate calls by making it possible to develop new labels for new referents. Thus, they are central to humans’ ability to articulate symbolic thought and coordinate unusually sophisticated levels of cooperation ^8^. However, few examples of learned vocal labeling are known in other species. Personal names are a type of vocal label that refer to another individual. Names must involve vocal learning, as an individual cannot be born knowing the names for all its future social affiliates. Thus, potential nonhuman analogs of personal names are highly relevant to understanding the evolution of language, and by extension, complex cognition and social behavior.

Most human words, including personal names, are arbitrary in structure; that is, they are not imitations of sounds typically made by the referent or tied to the physical properties of the referent ^3^. Arbitrariness is crucial to language because it enables communication about objects and ideas that do not make any imitable sound. However, clear evidence for arbitrary analogs of names in other species is lacking. Bottlenose dolphins (*Tursiops truncatus*) and some parrots (Psittacidae) address individual conspecifics by imitating the receiver’s “signature” call, a sound that is most commonly produced by the receiver to signal individual identity ^1,2,9^. When functioning as self-identification signals, these signature calls are indeed arbitrary ^10^. However, when other individuals copy a conspecific’s signature call to address them, it may be argued that the copied signature call is an iconic (non-arbitrary) label, since it is an imitation of a sound typically produced by the individual to whom the call refers. Non-imitative learned vocal labeling could allow communication about a wider range of referents than imitative labeling, but it may be more cognitively demanding, as it requires individuals to make an abstract connection between a sound and referent. Thus, if any non-human species were found to address individual conspecifics using labels that are not imitative of the receiver’s own calls, this would indicate a novel and perhaps uniquely complex form of communication with important implications for our understanding of language evolution and cognition.

Elephants are among the few mammals capable of mimicking novel sounds, although the function of this vocal learning ability is unknown ^11,12^. The most common call type produced by elephants is the rumble, a harmonically rich, low-frequency sound which is individually distinct ^13,14^, distinguishable, ^15^ and produced across most behavioral contexts ^16^. Contact rumbles (Supplementary Audio File S1) are long-distance calls produced when the caller is visually separated from one or more social affiliates and attempting to reinitiate contact, and greeting rumbles (Supplementary Audio File S2) are close-distance calls produced when one individual approaches another after a period of separation ^16^.

We analyzed contact and greeting rumbles from female-offspring groups of wild African savannah elephants to assess whether they contain individual vocal labels. We only used calls for which we were able to identify the caller and apparent intended receiver (527 calls from the greater Samburu ecosystem, northern Kenya, 98 from Amboseli National Park, southern Kenya). Receivers were identified as the individual who responded to the call by vocalizing or approaching the caller, the only adult member of the family group separated (>50m) from the caller when the caller produced a contact call, or the individual who approached/was approached by the caller when the caller produced a greeting call. We were able to determine which individuals were separated from the group at a given time by knowing the composition of each family group and by following the elephants for several hours each day and observing short-term fission and fusion events where some individuals split off from, lagged behind, and/or rejoined the rest of the group. Calls for which the receiver could not be identified or that appeared to be directed to multiple receivers (e.g., caller produced a contact call while separated from the whole family group) were excluded from analysis. We investigated (1) if elephants address conspecifics using receiver-specific vocal labels, (2) if the labels are imitative of the receiver’s calls or arbitrary, and (3) if different callers share the same label for the same receiver (Extended Data Table 1).

Our dataset consisted of 114 unique callers and 119 unique receivers, with 1-46 (median=2) calls per caller, 1-48 (median=2) calls per receiver, 1-9 (median=2) receivers per caller, and 1-10 (median=2) callers per receiver (Extended Data Fig. 1). For 597 of 625 calls, the caller and receiver belonged to the same family group. We measured two sets of acoustic features for each call (spectral and cepstral, see Supplementary Information; Extended Data Fig. 2) and ran all statistical models separately for each set of features. Results reported in the text and figures are for the spectral features only (see tables for results with cepstral features, which were similar).

### Calls were specific to individual receivers

We ran a random forest ^17^ with 6-fold cross-validation to predict the receiver of each of the 625 rumbles as a function of the acoustic features and compared the classification accuracy to a null distribution generated from 10,000 iterations of the same model with the acoustic features randomly permuted. We expected vocal labeling to only occur in contextually relevant calls, as humans and dolphins only use names or copied signature whistles in a minority of utterances ^18^. However, we used all 625 rumbles for analysis as there was no way to determine *a priori* which calls (or what proportion of calls) might contain a vocal label. Call structure varied clearly with the identity of the targeted receiver (Extended Data Fig. 3) as would be expected if elephants use vocal labels for other individuals. Our model correctly identified the receiver for 20.3% of calls analyzed, a significantly greater proportion than that of null models (permutation test, null models mean accuracy = 7.6 ± 0.75% correct, *P*<0.0001) (Fig. 1, Table 1), indicating receivers of calls could be correctly identified from call structure statistically significantly better than chance.

**Figure 1.**
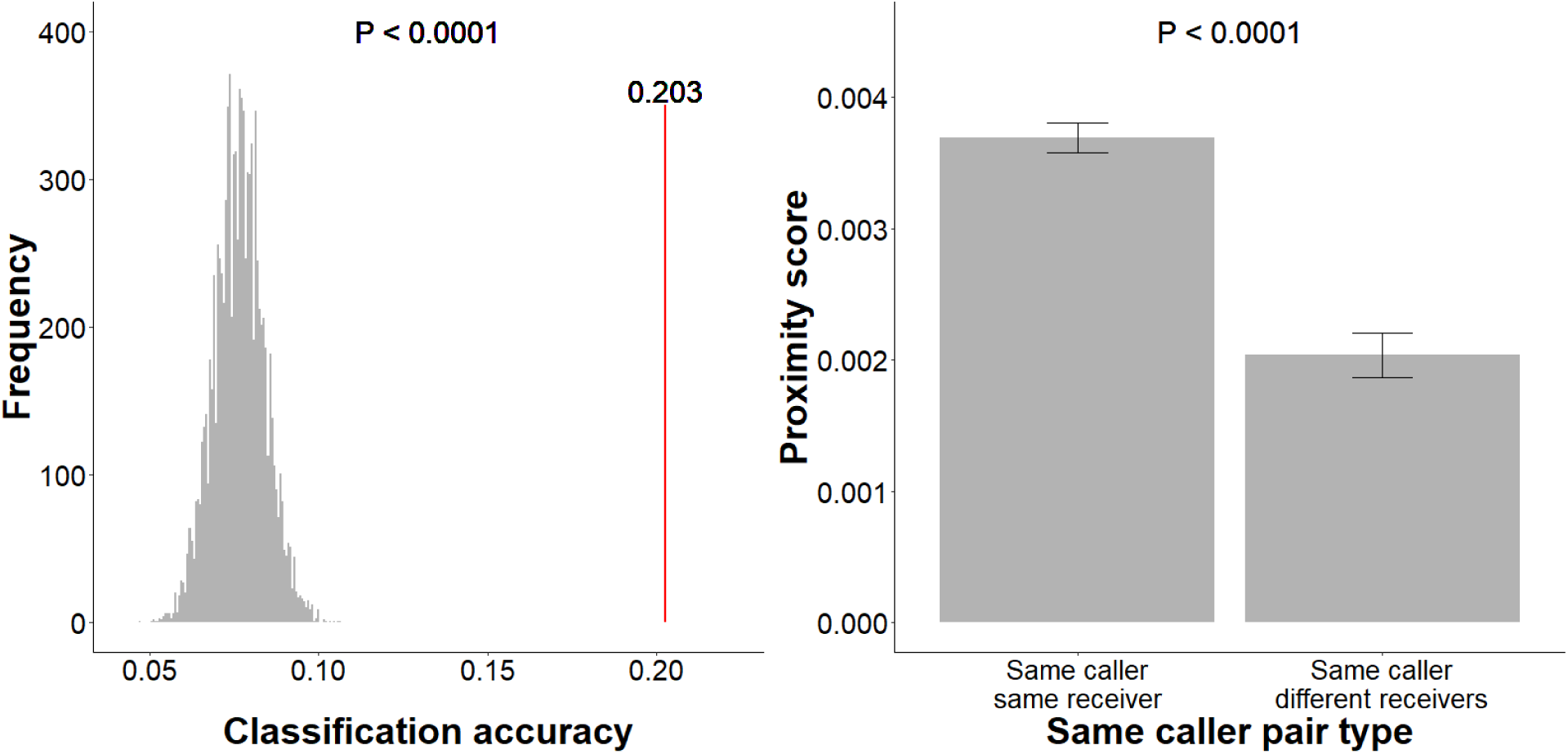
Evidence that calls are specific to individual receivers within a caller. **Left:** classification accuracy of random forest predicting receiver ID from acoustic features (red line) was significantly higher than classification accuracies of 10,000 null models predicting receiver ID from randomized acoustic features (gray histogram). Cross-validation folds were stratified so that model was trained and test on same combinations of caller and receiver; thus, classification accuracy represents receiver specificity of calls within a caller. **Right:** calls with the same caller and same receiver were significantly more similar (higher proximity score) than calls with the same caller and different receivers (ANOVA on ranks). Error bars represent standard errors of the mean.

**Table 1.**
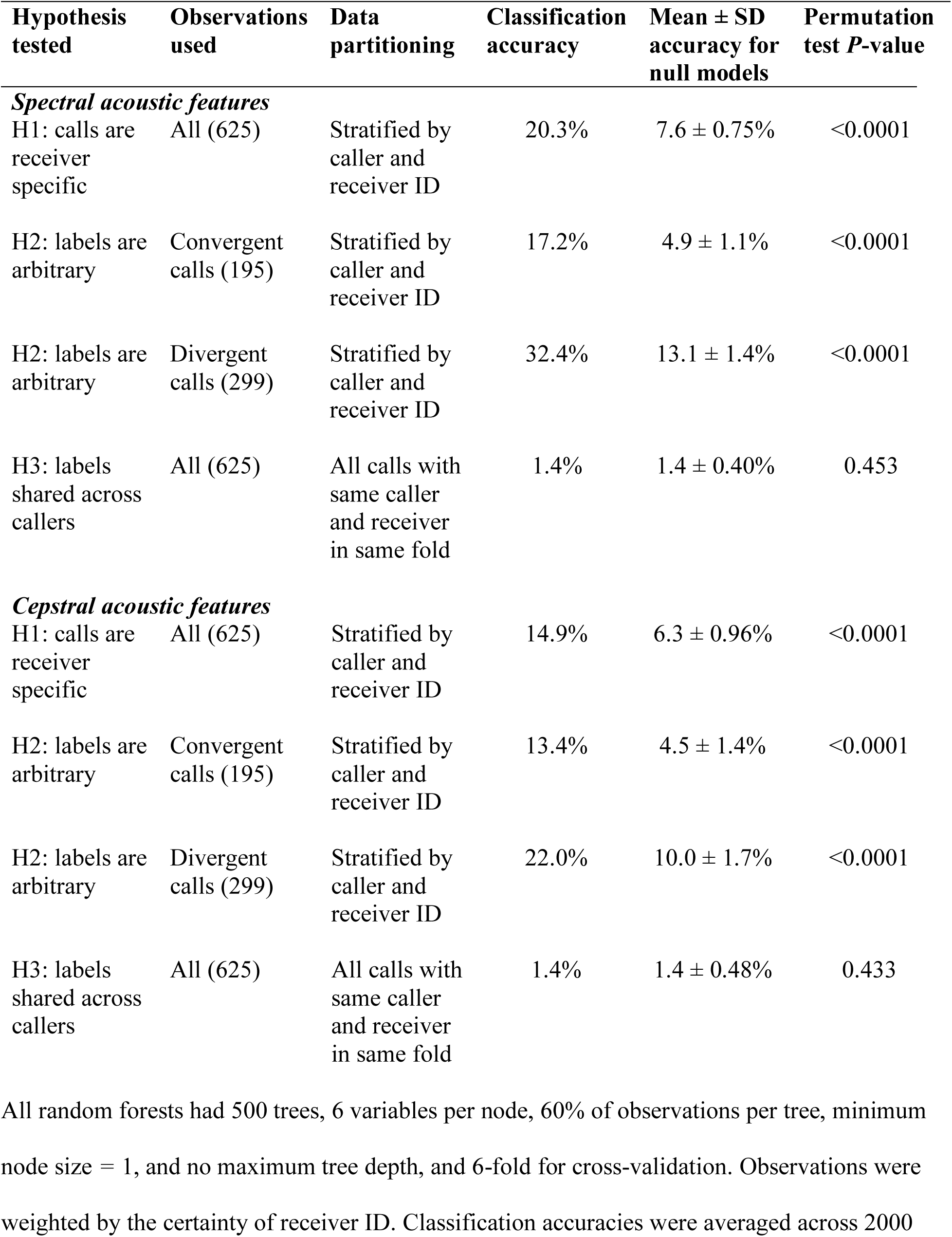
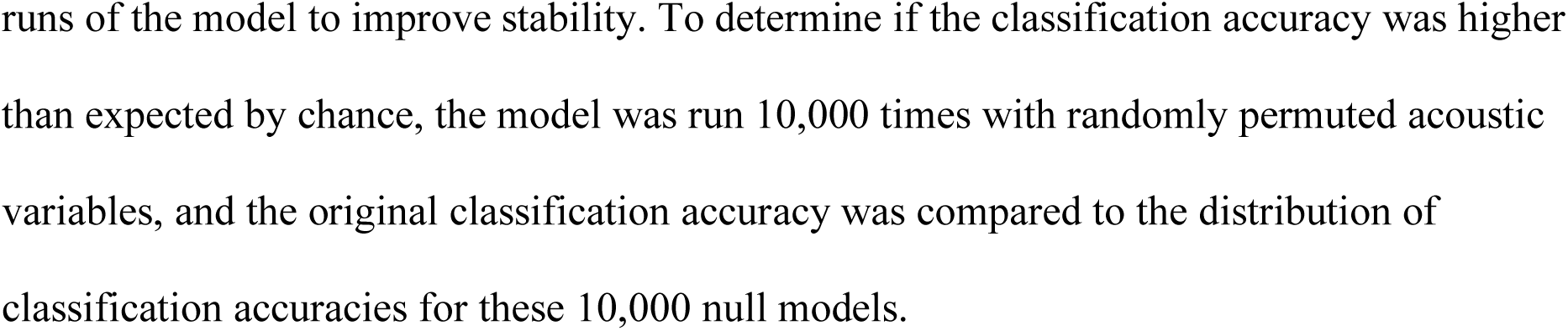
Results of random forest models predicting receiver ID as a function of the acoustic features.

| Hypothesis tested | Observations used | Data partitioning | Classification accuracy | Mean ± SD accuracy for null models | Permutation test $P$-value |
| --- | --- | --- | --- | --- | --- |
| *Spectral acoustic features* |  |  |  |  |  |
| H1: calls are receiver specific | All (625) | Stratified by caller and receiver ID | 20.3% | $7.6 \pm 0.75\%$ | <0.0001 |
| H2: labels are arbitrary | Convergent calls (195) | Stratified by caller and receiver ID | 17.2% | $4.9 \pm 1.1\%$ | <0.0001 |
| H2: labels are arbitrary | Divergent calls (299) | Stratified by caller and receiver ID | 32.4% | $13.1 \pm 1.4\%$ | <0.0001 |
| H3: labels shared across callers | All (625) | All calls with same caller and receiver in same fold | 1.4% | $1.4 \pm 0.40\%$ | 0.453 |
| *Cepstral acoustic features* |  |  |  |  |  |
| H1: calls are receiver specific | All (625) | Stratified by caller and receiver ID | 14.9% | $6.3 \pm 0.96\%$ | <0.0001 |
| H2: labels are arbitrary | Convergent calls (195) | Stratified by caller and receiver ID | 13.4% | $4.5 \pm 1.4\%$ | <0.0001 |
| H2: labels are arbitrary | Divergent calls (299) | Stratified by caller and receiver ID | 22.0% | $10.0 \pm 1.7\%$ | <0.0001 |
| H3: labels shared across callers | All (625) | All calls with same caller and receiver in same fold | 1.4% | $1.4 \pm 0.48\%$ | 0.433 |

All random forests had 500 trees, 6 variables per node, 60% of observations per tree, minimum node size = 1, and no maximum tree depth, and 6-fold for cross-validation. Observations were weighted by the certainty of receiver ID. Classification accuracies were averaged across 2000
runs of the model to improve stability. To determine if the classification accuracy was higher than expected by chance, the model was run 10,000 times with randomly permuted acoustic variables, and the original classification accuracy was compared to the distribution of classification accuracies for these 10,000 null models.

To determine if this could be an artifact of the correlation between caller ID and receiver ID in our dataset, we controlled for caller ID by comparing the mean similarity of pairs of calls with the same caller and receiver to the mean similarity of pairs of calls with the same caller and different receivers, using proximity scores derived from the random forest as a metric of call similarity ^19^. To control for the possibility that calls were specific to the type of relationship between the caller and receiver rather than to the individual receiver per se, we categorized social relationship based on relatedness and age (a proxy for dominance) (Extended Data Table 3), and only considered pairs of calls with the same type of relationship between caller and receiver. Calls with the same caller and same receiver were significantly more similar than calls with the same caller and different receivers, even after controlling for social relationship, behavioral context, and recording date, further supporting the hypothesis that rumbles are specific to individual receivers (ANOVA, *F*_1_=94.61, *P*<0.0001, Cohen’s D=0.412) (Fig. 1, Extended Data Table 4). As calls in our dataset were predominantly between individuals in the same family group, our results only provide evidence for vocal labeling within family groups.

### Vocal labelling likely does not rely on imitation of receiver

If calls are imitative of the receiver’s calls, then callers should sound more like a given receiver when addressing her than when addressing other individuals. Pairs of calls in which the receiver of one call was the caller of the other call were slightly but significantly more similar on average than pairs in which this was not the case, suggesting possible imitation of the receiver’s calls (ANOVA, *F*_1_=11.70, *P*=0.0006, Cohen’s D=0.0037) (Extended Data Table 5). However, given the exceedingly small effect size (0.78% of SD) and large sample size of call pairs (n=11,309), this significant difference may not be biologically meaningful. Moreover, among the calls for which we had recordings of the receiver and recordings of the caller addressing other individuals (n=494), 60.5% were divergent from the receiver’s calls; that is, less similar to the receiver’s calls than typical for that caller (see Supplementary Information). The classificatory model performed significantly better than the null model for both convergent and divergent calls (convergent calls: 17.2% correct, null models mean accuracy = 4.9 ± 1.1%, *P*<0.0001; divergent calls: 32.4% correct, null models mean accuracy = 13.1 ± 1.4%, *P*<0.0001) (Fig. 2, Table 1). Finally, among both convergent and divergent calls, calls with the same caller and same receiver were more similar than calls with the same caller and different receivers (ANOVA; convergent calls: *F*_1_=15.30, *P*=0.0001, Cohen’s D=0.411; divergent calls: *F*_1_=8.67, *P*=0.0033, Cohen’s D=0.262) (Fig. 2, Extended Data Table 4). This suggests that vocal labeling in elephants likely does not rely on imitation of the receiver’s calls. While we cannot rule out the possibility that elephants imitated calls made by the receiver that were not included in our dataset, elephants are not known to produce discrete “signature” calls like dolphins and parrots; instead, the caller-specificity of elephant rumbles is likely a product of voice characteristics that are present across calls ^13,14^.

**Figure 2.**
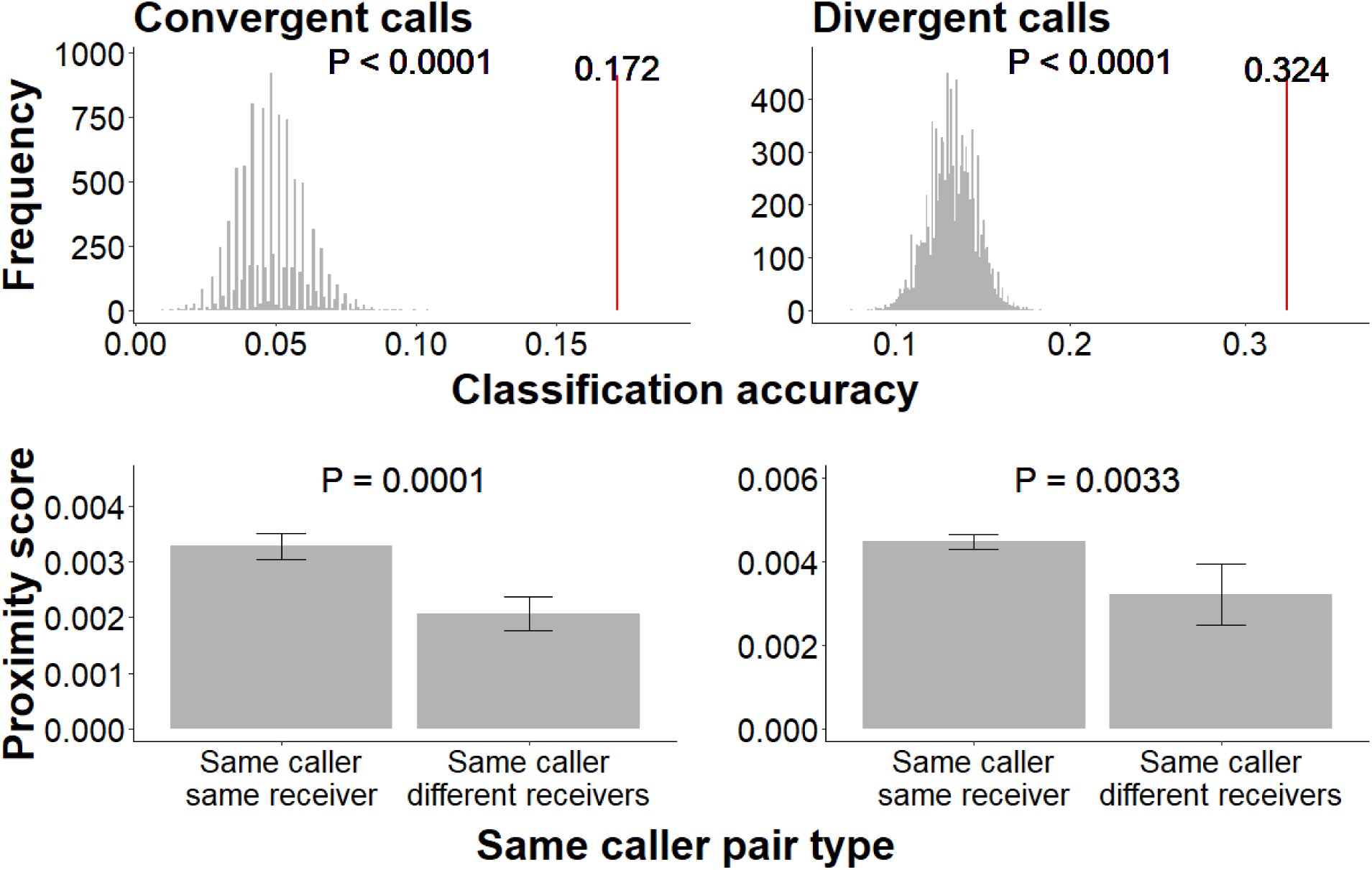
Evidence that vocal labeling likely did not rely on imitation of the receiver’s calls. Random forest predicted receiver ID significantly better than null models both among calls that were identified as convergent to receiver’s calls (**top left**) and divergent from receiver’s calls (**top right**). Pairs of calls with the same caller and same receiver were more similar (higher proximity score) than pairs of calls with the same caller and different receivers, both among calls that were convergent to receiver’s calls (**bottom left**) and calls that were divergent from receiver’s calls (**bottom right**) (ANOVA on ranks). In top row, red lines represent classification accuracy of original random forest model and gray histograms represent distribution of classification accuracies of null models with randomized acoustic features. In bottom row, error bars represent standard errors of the mean.

### Mixed evidence for convergence among callers addressing same receiver

In humans and bottlenose dolphins, different callers generally use the same label for a given receiver. To determine if different callers use similar labels to address the same receiver in elephants, we ran a random forest structured to predict receiver ID from different callers than the model was trained on. This model correctly classified 1.4% of calls, no better than the corresponding null models (permutation test, mean accuracy of null models=1.4 ± 0.40% correct, *P=*0.453) (Fig. 3, Table 1). However, calls from different callers to the same receiver were significantly more similar on average than calls from different callers to different receivers (ANOVA, *P*<0.0001, Cohen’s D=0.134) (Fig. 3, Extended Data Table 6). These mixed results may be due to the fact that rumbles simultaneously encode multiple messages ^13,16,20,21^. If vocal labels account for only a small portion of the variation in rumbles, the random forest may have been influenced by context or caller-specific features, thus reducing its ability to predict receiver ID across callers, even if different callers address the same receiver with the same label. Further work to identify how vocal labels are encoded in elephant calls will be necessary to definitively determine if different callers use the same label for the same receiver.

**Figure 3.**
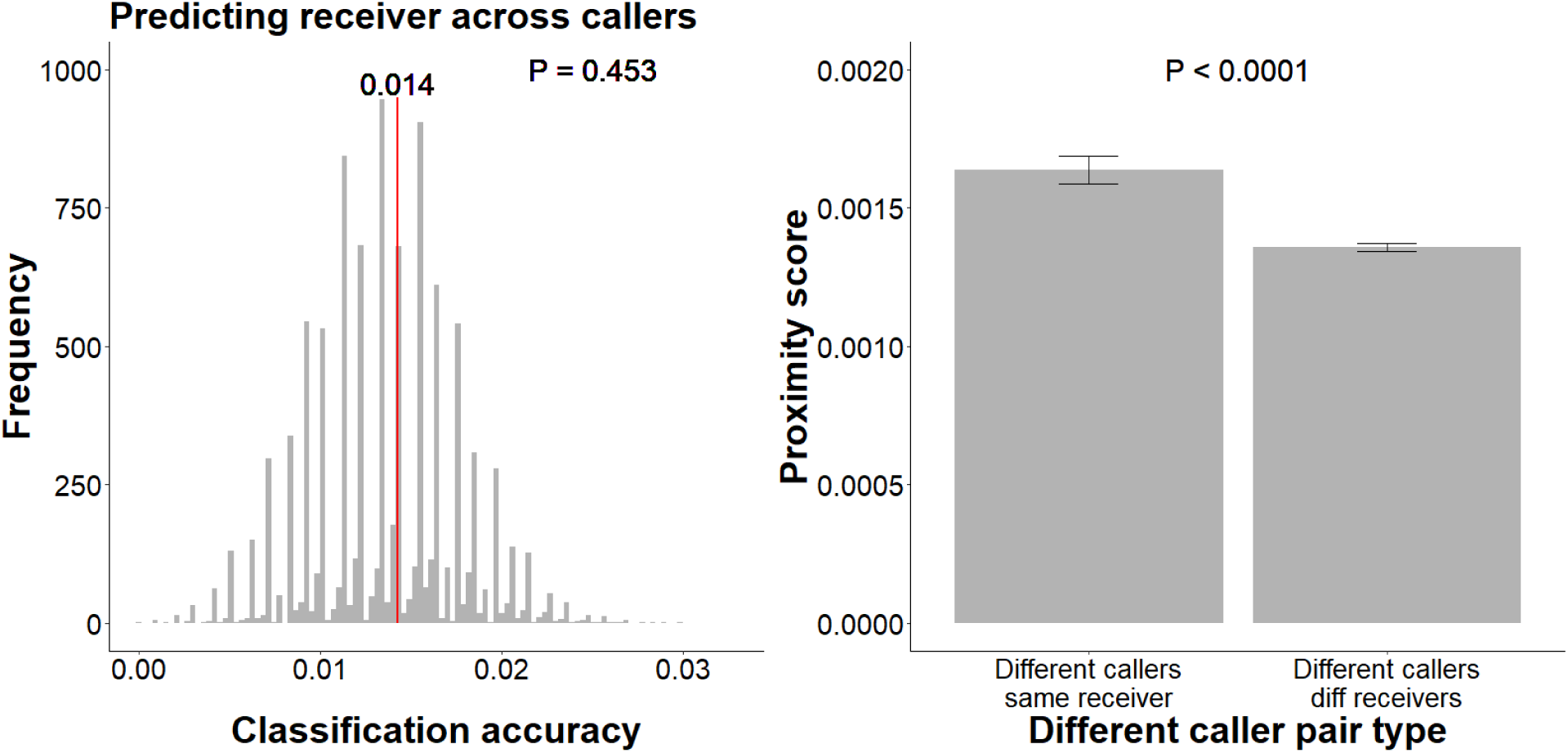
Mixed evidence that different callers use similar labels for the same receiver. **Left:** Classification accuracy (red line) of random forest designed to predict receiver ID from acoustic features independently of caller ID (all calls with the same caller and receiver allocated to the same cross-validation fold) was not significantly different from classification accuracies of null models with randomized acoustic features (gray histogram). **Right:** Pairs of calls with different callers and the same receiver were significantly more similar (higher proximity score) than pairs of calls with different callers and different receivers (ANOVA on ranks). Error bars represent standard errors of the mean.

### Elephants responded more strongly to playback of calls originally addressed to them

To determine if elephants perceive and respond to the vocal labels in calls addressed to them, we compared reactions of 17 wild elephants to playback of a call that was originally addressed to them (test) relative to playback of a call from the same caller that was originally addressed to a different individual (control). By using test and control stimuli from the same caller, we controlled for the possibility of the caller’s relationship to the subject influencing the results. To control for the possibility that calls are specific to the type of relationship between the caller and receiver rather than to the individual receiver per se, we included the type of relationship between the caller and the original receiver of the call as a factor in the analysis.

Further supporting the existence of vocal labels, subjects approached the speaker more quickly (Cox regression, χ^2^*=*6.8, *P=*0.009) and vocalized more quickly (Cox regression, χ^2^*=*7.9, *P=*0.005) in response to test playbacks than control playbacks (Fig. 4, Table 2). They also produced more vocalizations in response to test playbacks, although this model failed to converge (Poisson regression, χ^2^*=*6.2, *P=*0.013) (Fig. 4, Table 2). There was no significant difference between test and control trials in latency to vigilance (Cox regression, χ^2^*=*3.1, *P=*0.08) or in the change in vigilance duration before and after the playback (linear regression, χ^2^*=*0.06, *P=*0.81), although there was a nonsignificant trend toward faster onset of vigilance in test trials (Table 2).

**Figure 4.**
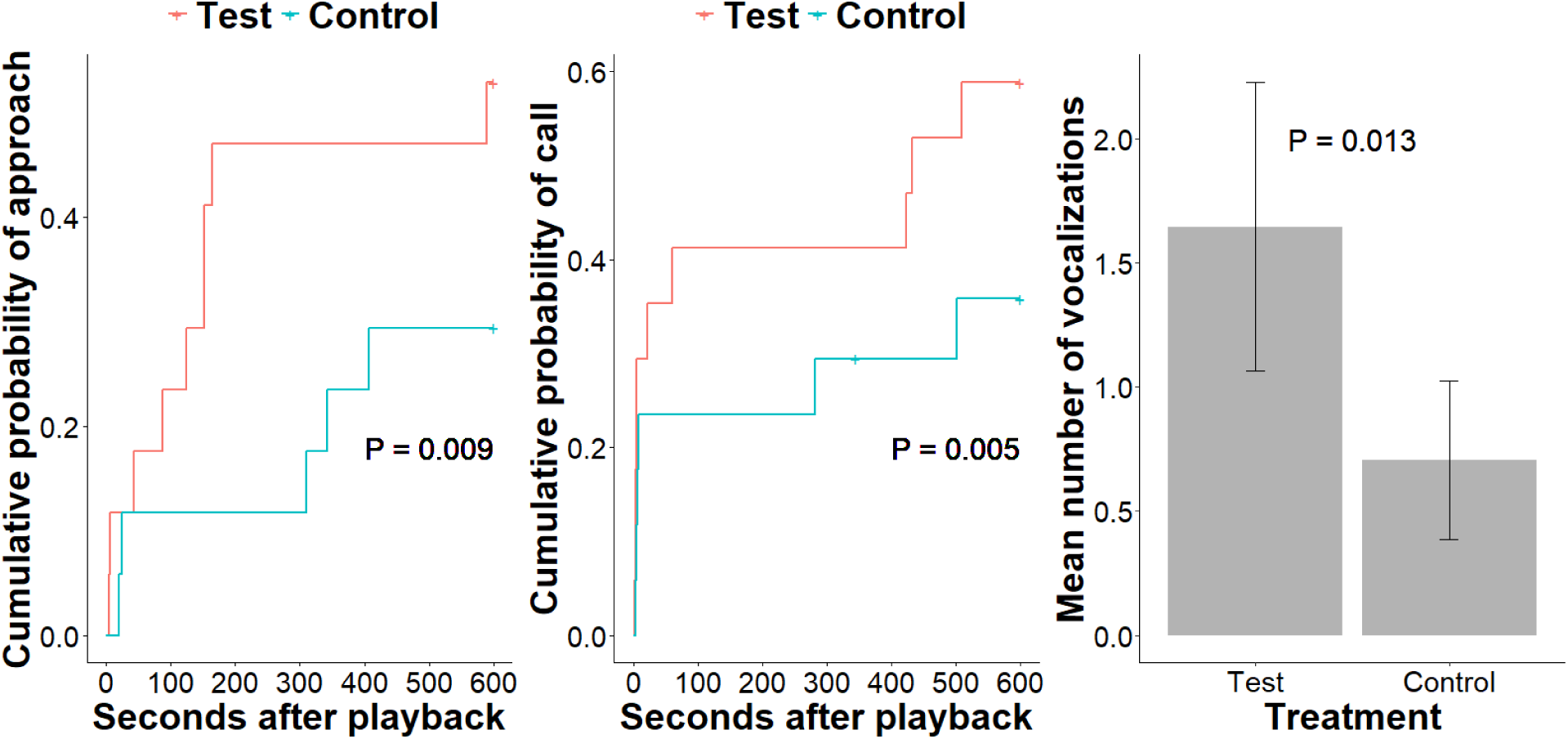
Response to playbacks of test stimuli (calls originally addressed to the subject) vs. control stimuli (calls from the same caller originally addressed to a different individual). Subjects approached the speaker more quickly (**left;** Cox regression), vocalized more quickly (**center;** Cox regression), and produced more vocalizations (**right;** Poisson GLM) in response to test playbacks than controls (note the model for number of vocalizations failed to converge). Error bars in rightmost panel represent standard errors of the mean.

**Table 2.**
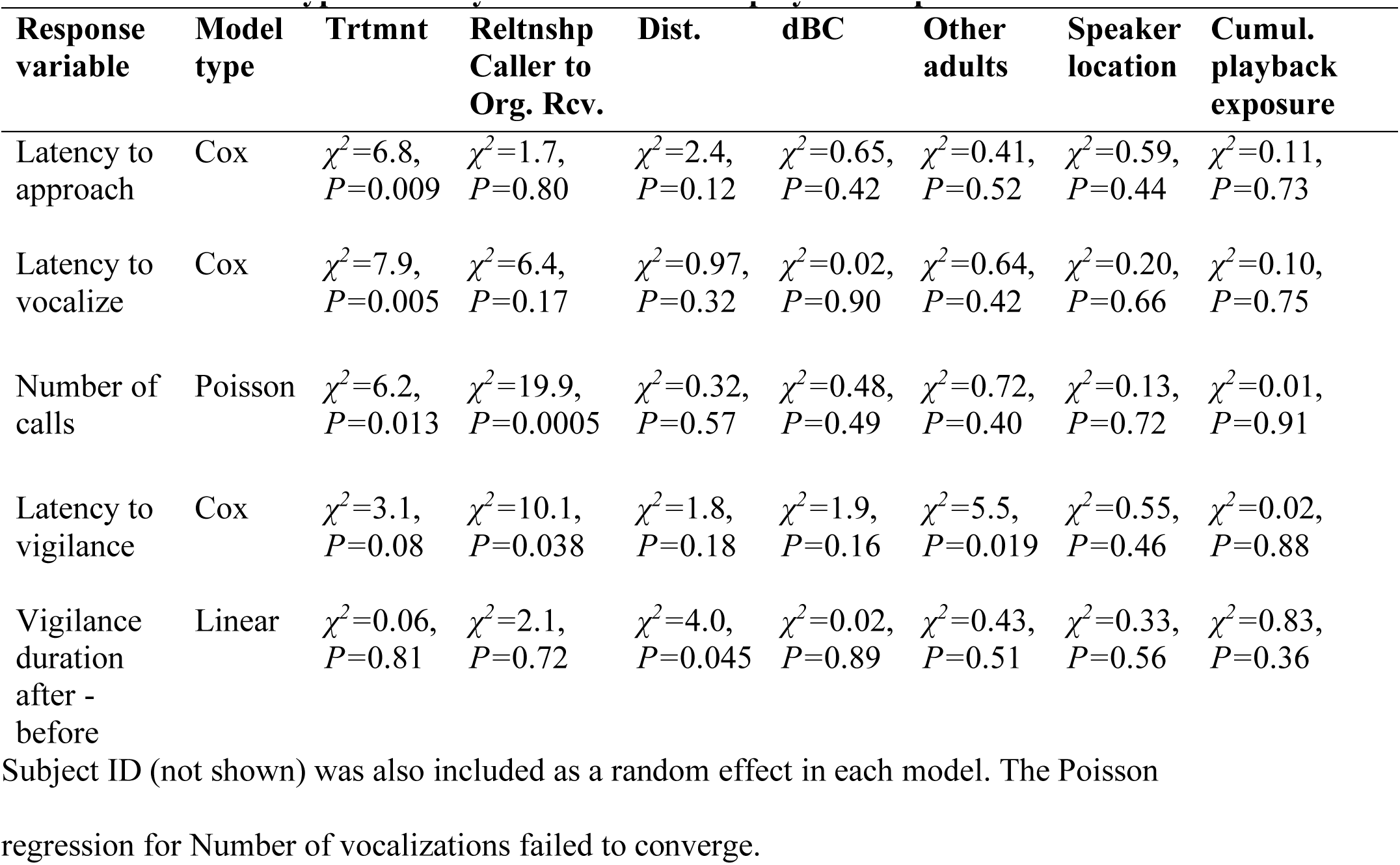
Results for Type III Analyses of Deviance on playback experiment models.

## Discussion

To our knowledge, this study presents the first evidence for vocal addressing of conspecifics without imitation of the receiver’s calls in nonhuman animals. Very few species are known to address conspecifics with vocal labels of any kind. Where evidence for vocal labels has been found, they are either clearly imitative ^1,2,9^ or of unknown structure ^22,23^. Our data suggest that elephants label conspecifics without relying on imitation of the receiver’s calls, a phenomenon previously known to occur only in human language.

The social behavior and ecology of elephants create an environment in which individual vocal labeling may be particularly advantageous. Due to their fission-fusion social dynamics, elephants are often out of sight of their closely bonded social partners and produce contact rumbles to communicate over long distances ^16,24^. Characteristic fission-fusion dynamics in elephants include coordinated movement to and from resources while proximately diffusing to avoid foraging competition ^25,26^. Vocal labels could enhance coordinating ability when out of sight of one another. In contact calling scenarios, vocal labeling could allow callers to attract the attention of a specific intended receiver. While greeting rumbles are produced in close proximity when the caller and receiver typically have visual contact ^16^, vocal labeling in greeting rumbles could possibly strengthen social bonds with specific individuals. Humans experience a positive affective response and increased willingness to comply with requests when someone remembers their name ^27^.

Nonetheless, the fact that our random forest model correctly predicted receiver ID for only around a fifth of calls (albeit significantly better than random) suggests that vocal labels only occur in a minority of rumbles and thus are likely not necessary in all or even most contexts. For example, contact and greeting calls may occur in vocal sequences where labeling the receiver in each call would be redundant ^16^, and in the dry season, when elephant families fission into smaller groups, there may be less ambiguity about the intended receiver in many scenarios ^26^. Indeed, both humans and bottlenose dolphins only use individual vocal labels (i.e., names or imitated signature whistles) in a small percentage of utterances ^18^.

When vocal labels do occur, they are likely only one component among many in the call. Rumbles are recognized to simultaneously encode multiple messages, including but not limited to caller identity, age, sex, emotional state, and behavioral context ^13,16,20,21^. Moreover, the top acoustic features for predicting receiver ID were not those that explained the most variation in the calls (see Supplementary Information), suggesting that vocal labels account for only a small fraction of the total variation in rumbles. Rather than comprising a whole stand-alone call, elephant vocal labels may be embedded within a call that simultaneously conveys multiple additional messages. The richness in the information content of elephant vocalizations makes it difficult to identify the specific acoustic parameters that encode receiver ID. Unlike dolphin signature whistles ^18^, elephant vocal labels cannot be discerned by visual inspection of the spectrogram and are likely encoded by a complex and subtle interaction among many acoustic parameters. As a result, we employed machine learning in this analysis, but innovative approaches in signal processing may be necessary to isolate the vocal labels within rumbles.

Both African and Asian elephants have a demonstrated capacity for vocal mimicry in captivity, but no prior study has documented a function of this ability in the wild ^11,12^. We speculate that vocal labeling may be one, if not the primary, function of vocal production learning in wild elephants. Dolphins and parrots, which show evidence for individual vocal labeling via imitation of the receiver, are adept vocal learners. Another vocal learner, the Egyptian fruit bat (*Rousettus aegyptiacus*), produces calls that are specific to individual receivers and may be vocal labels as well, although it is currently unknown if the bats perceive this information ^23^. Taken together, this raises the possibility that social selection pressures creating a need to address individual conspecifics may have led to multiple independent origins of vocal production learning.

The use of learned arbitrary labels is part of what gives human language its uniquely broad range of expression ^3^. Our results suggesting that wild elephants also use arbitrary vocal labels for individual conspecifics provide an opportunity to investigate the selection pressures that may have led to the evolution of this rare ability in two divergent lineages. Moreover, these findings raise intriguing questions about the complexity of elephant social cognition, considering the potential relevance of symbolic communication to their social decision making.

## METHODS

### Field recording

We collected audio recordings of wild female-calf groups in Amboseli National Park, Kenya in 1986-1990 and 1997-2006 and Samburu and Buffalo Springs National Reserves (hereafter, Samburu), Kenya in Nov 2019-Mar 2020 and Jun 2021-Apr 2022. Both populations have been continuously monitored for decades and all individuals can be individually identified by external ear morphology ^26,28^. We recorded calls from a vehicle during daylight hours with all-occurrence sampling ^29^ using an Earthworks QTC1 microphone (4 Hz-40 kHz ± 1 dB) with a Nagra IV-SJ reel-to-reel tape recorder or an HHB PDR 1000 DAT recorder in Amboseli, and an Earthworks QTC40 microphone (3 Hz-40kHz ± 1 dB) with a Sound Devices MixPre3 or MixPre3-II digital recorder in Samburu. Recordings were recorded at a 48 kHz sampling rate with 16 bits of amplitude resolution and stored at 2 kHz in Amboseli and recorded and stored at 44.1 kHz with 24 or 32 bits of amplitude resolution in Samburu.

When possible, we recorded for each call the identity of the caller, the behavioral context, and the identity of the receiver (criteria for identifying receiver defined in Main Text). The caller was identified using behavioral and contextual cues, such as an open mouth, flapping ears, or being the only individual of the right age class in the immediate vicinity ^16^. We scored behavioral context according to a published methodology ^16^. For each call, we recorded the certainty with which we knew the caller ID, behavioral context, and receiver ID as a number between 0 and 1 (see Supplementary Information). In all statistical analyses, we weighted each call by the certainty of receiver ID, so calls with greater certainty about the identity of the receiver would have a proportionally greater impact on the model.

### Acoustic analysis

We only included in analysis contact and greeting rumbles with certainty of caller ID, receiver ID, and behavioral context greater than 0, with no significant overlap with other calls or other loud sounds in the same frequency range, and that were recorded close enough to the microphone for the first two formants to be clearly visible in the spectrogram (98 rumbles from Amboseli, 527 from Samburu). We performed all acoustic and statistical analyses in R version 4.1.3 ^30^. We automatically detected the onset and offset of each call from the amplitude envelope using the function segment() in the package soundgen ^31^, manually adjusting the detected times when necessary. We then measured two alternative sets of features: spectral and cepstral (see Supplementary Information). The spectral features consisted of the smoothed Hilbert amplitude envelope (350 ms moving average window, 90% overlap), the vectors of energy values in 26 mel-frequency bands between 0-500 Hz (measured at 35 ms intervals), and the vectors of delta and delta-delta coefficients for the mel-frequency bands (79 vectors total) (Extended Data Fig. 2). The cepstral features consisted of the amplitude envelope, the vectors of the first 12 mel-frequency cepstral coefficients measured at 35 ms intervals, and the vectors of delta and delta-delta coefficients for the cepstral coefficients.

As the raw acoustic vectors (mel spectral bands, MFCCs, and their delta and delta-delta values) represented a matrix of values for each call, it was necessary to calculate lower-dimensional derived features from these matrices as input variables for statistical models. We calculated derived features separately for the spectral and cepstral features. In brief, we scaled the acoustic vectors and decorrelated them with a robust principal components analysis using the rpca package in R, which decomposes the data into a robust matrix and a sparse matrix containing the outlier values (λ=0.00996) ^32^. The final derived features we calculated were the median, robust skewness, minimum extent, and equivalent statistical extent of the sparse matrix, the means of the first *n* low-rank principal components required to explain 99.9% of the variation (74 for spectral features, 12 for cepstral features), and 8 measures of the spectral properties of the low-rank principal components, calculated by treating each principal component as if it were a waveform (see Supplementary Information) (Extended Data Table 2).

### Statistical analysis of acoustic data

#### Are calls specific to individual receivers (hypothesis 1)?

We ran a 6-fold cross-validated random forest model in the R package ranger ^33^ to predict the identity of the receiver of each call (receiver ID) as a function of the acoustic features. We stratified the cross-validation folds by caller ID and receiver ID to ensure as even a distribution as possible of all caller-receiver dyads across all folds. Thus, if calls contain acoustic cues to receiver ID, this model was expected to predict receiver ID better than chance regardless of whether the label for a given receiver is shared across callers (Extended Data Table 1, hypothesis 1, prediction 1). The model used 625 observations, 500 trees, 6 variables per node, 60% of observations per tree, a minimum node size of 1, and no maximum tree depth, and observations were weighted by certainty of receiver ID. To increase the stability of the model’s classification accuracy, we ran the model 2000 times and used the mean classification accuracy across the 2000 runs. To determine if the model predicted receiver ID better than expected by chance, we ran the model 10,000 times with the acoustic features randomly permuted and compared the classification accuracy of the original model (averaged across 2000 runs) to the null distribution of classification accuracies generated by the 10,000 models with randomized acoustic features.

As caller ID and receiver ID were partially aliased in our dataset (Extended Data Fig. 1), the random forest could theoretically use acoustic cues to caller ID ^16^ to predict receiver ID, even if the calls did not contain any vocal label identifying the intended receiver. To disentangle the effects of caller ID and receiver ID on call structure, we compared the mean pairwise similarities between pairs of calls with the same caller and receiver and pairs with the same caller and different receivers (Same Caller Pair Type). As a metric of call similarity, we extracted a proximity score for each pairwise combination of calls from a random forest trained to predict receiver ID as a function of the acoustic features on the full dataset (625 training observations, 8000 trees, other hyperparameters and weighting same as above). The proximity score for a given pair of calls was the proportion of trees in which both calls were classified in the same terminal node, corrected for the size of each node, and represented the degree of similarity between the two calls in terms of the acoustic features most relevant to predicting receiver ID ^19^. If calls are specific to individual receivers within a given caller, then pairs of calls with the same caller and same receiver should be more similar (have higher proximity scores) than pairs of calls with the same caller and different receivers (Extended Data Table 1, hypothesis 1, prediction 2).

Previous work has shown that elephants vary the structure of their rumbles when interacting with more dominant vs. more subordinate conspecifics ^13^. To rule out the possibility that calls were specific to the type of relationship between caller and receiver rather than to individual receivers *per se*, we restricted the analysis of Same Caller Pair Type to pairs of calls that had the same type of relationship between caller and receiver. We defined caller-receiver relationship using 12 categories based on sex, family group membership, relative age, and mother-offspring relationship, reflecting the fact that dominance in elephants is primarily determined by age ^34,35^ and that mother-calf bonds are the strongest social bonds in elephants ^26,36^ (Extended Data Table 3). We also excluded pairs of calls that were recorded on the same date, as preliminary analyses indicated that calls recorded on the same day were more similar than calls recorded on different days, likely due to similarities in ambient conditions and/or autocorrelation within a calling bout (final sample size = 2391 call pairs). As calls from different behavioral contexts differ in acoustic structure ^16^, we categorized each pair of calls according to whether the two calls had the same or different behavioral contexts (“Same Context”) and included this variable as a factor in the analysis.

The proximity scores were highly skewed to the right, so we rank-transformed them and ran a Type III ANOVA with rank-transformed proximity score as the response variable and Same Caller Pair Type and Same Context as the factors. We weighted each observation (pair of calls) in the model by the minimum value of the certainty of caller ID and certainty of receiver ID for the two calls in the pair.

#### Are vocal labels based on imitation of the receiver’s calls (hypothesis 2)?

If elephants imitate the calls of the receiver that they are addressing, then callers should sound more like a given conspecific when they are addressing her than when they are addressing someone else (Extended Data Table 1, hypothesis 2, prediction 1). To assess whether this was the case, we classified each pair of calls into one of two types (hereafter, “Imitation Pair Type”): pairs in which the receiver of one call was the caller of the other call, and pairs in which this was not the case. We separately classified each call pair according to whether the two calls had the same relationship between caller and receiver (hereafter, “Same Relationship”). We also created a categorical variable Caller Dyad ID, which was an identifier for each unique combination of callers that comprised a call pair. We ran a Type III ANOVA with rank-transformed proximity score as the response variable and Imitation Pair Type, Same Relationship, Same Context, and Caller Dyad ID as factors. We weighted each observation (pair of calls) in the model by the minimum value of the certainty of caller ID and certainty of receiver ID for the two calls in the pair. By controlling for Caller Dyad ID in the model we assessed the effect of Imitation Pair Type within a given pair of callers; that is, whether calls from caller A to receiver B were more similar to the receiver B’s calls than calls from the caller A addressed to other receivers were to receiver B’s calls. Pairs of calls that had the same caller or receiver, were recorded on the same day, were recorded from different family groups, or for which Caller Dyad ID did not occur with both levels of Imitation Pair Type were excluded from analysis (final sample size = 11,309 call pairs). Pairs of calls from different family groups were excluded because they comprised a small percentage of pairs where the receiver of one call was the caller of the other, and because it is possible that different families have different “dialects” which would influence call similarity.

If vocal imitation of the receiver occurs, it might or might not be the mechanism behind individual vocal labeling. To assess whether imitation of the receiver’s calls was necessary for vocal labeling, we examined the calls in the dataset for which we had at least one recording of the receiver’s calls and at least one recording of the caller addressing someone other than the receiver (n=494). For each of these calls, we calculated the mean proximity score between the focal call and all the calls made by the receiver (Mean Proximity to Focal Receiver When Targeting Focal Receiver) as well as the mean proximity score between each of the calls made by the focal caller to an individual other than the focal receiver and each of the calls made by the focal receiver (Mean Proximity to Focal Receiver When Targeting Others). Calls in which the Mean Proximity to Focal Receiver When Targeting Focal Receiver was greater than the Mean Proximity to Focal Receiver When Targeting Others were classified as “convergent” (n=195) and divergent otherwise (n=299). We then examined the proportion of convergent and divergent calls that were classified correctly by the random forest model with receiver ID and the acoustic features as input variables, and cross-validation folds stratified by caller ID and receiver ID. If vocal labeling relies on imitation of the receiver’s calls, we expected only the convergent calls to be classified correctly more often than by the null model, but if imitation is not necessary for vocal labeling, we expected both convergent and divergent calls to be classified correctly more often than by the null model (Extended Data Table 1, hypothesis 2, prediction 2). We also ran separate ANOVAs for the convergent calls and divergent calls, with rank-transformed proximity score as the response and Same Caller Pair Type and Same Context as the factors (excluding pairs of calls recorded on the same day). If vocal labeling relies on imitation of the receiver, we expected that there would only be an effect of Same Caller Pair Type among the convergent calls, but if imitation is not necessary for vocal labeling, we expected to observe an effect of Same Caller Pair Type among both sets of calls (Extended Data Table 1, hypothesis 2, prediction 3).

#### Do different callers use the same label for the same receiver (hypothesis 3)?

To determine if different callers use the same label for the same receiver, we ran another 6-fold cross-validated random forest model to predict receiver ID as a function of the acoustic features but partitioned the cross-validation folds such that all calls with the same caller and receiver were always allocated to the same fold (hyperparameters and weighting same as first model). This model tested whether receiver ID could be predicted independently of caller ID, which should only be possible if different callers use similar labels for a given receiver (Extended Data Table 1, hypothesis 3, prediction 1). We averaged the classification accuracy of the model across 2000 runs and compared this value to the distribution of classification accuracies generated by 10,000 iterations of the same model with the acoustic features randomly permuted.

If different callers use similar labels for the same receiver, then pairs of calls with different callers and the same receivers should be more similar than pairs of calls with different callers and different receivers (Extended Data Table 1, hypothesis 3, prediction 2). To test whether this was the case, we ran another Type III ANOVA with rank-transformed proximity score as the response variable and Different Caller Pair Type (different callers/same receiver or different callers/same receiver), Same Relationship, and Same Context as the factors. As before, we excluded pairs of calls recorded on the same date or from different family groups (final sample size = 20,235 call pairs).

#### How are labels encoded in calls?

To investigate which acoustic features encode receiver ID and caller ID we extracted variable importance scores (Supplementary Table S1) from a conditional inference random forest model in the R package “party” ^37^ trained on the full dataset to predict the response variable in question (receiver ID or caller ID) as a function of the acoustic features and weighted by the certainty of the response variable (625 training observations, 8000 trees, all other hyperparameters same as other random forests). We used a conditional inference forest because unlike traditional random forest, it is not biased towards correlated variables ^37^. We only calculated variable importance scores for the spectral features, as cepstral coefficients are difficult to interpret intuitively. To assess the relative importance of the original acoustic contours, we weighted the loadings of the acoustic contours on each principal component by the variable importance score of the mean of the principal component in question, and then calculated the sum of the absolute values of these weighted loadings for each acoustic contour (Supplementary Table S2). Acoustic contours with a higher sum of the absolute values of the weighted loadings were deemed more important. This weighting process only considered the means of low-rank principal components, as it was not clear how to relate the other features back to the original acoustic contours. However, means of low-rank principal components accounted for the top 19 variables for the receiver ID model and top 33 variables for the caller ID model.

### Playback experimental design

To determine if elephants respond more strongly to calls addressed to them (Extended Data Table 1, hypothesis 1, prediction 3), we played back rumbles with known adult female callers and known receivers to 17 elephants (15 adult females, one 9yo female, one 9-10yo male) in the Samburu study area. Fourteen subjects received one “test” playback of a call that was originally addressed to them and one “control” playback of a call from the same caller that was originally addressed to another individual. One subject received two sets of test and control playbacks from two different callers, one received only a test playback, and one received only a control playback (Extended Data Table 7). Most stimuli functioned as the test stimulus for one subject and the control stimulus for another, but no stimulus was used as the same experimental condition for more than one subject. Order of presentation was balanced across subjects, and we waited at least 7 days (mean *=* 29.5 ± 27.1 days) between successive playbacks to the same subject.

### Playback stimuli

Playback stimuli were recorded in Samburu and Buffalo Springs between January 2020 and March 2022 from adult female callers. In all but two cases, the playback stimuli were contact calls. In one case we used a loud greeting call because we were unable to record a contact call from the caller in question, and in one case we used a call that was produced in a similar context to contact calls (caller and receiver >100 m apart and out of sight of each other), but was lower in amplitude than a typical contact call and was part of a lengthy antiphonal exchange between two individuals, and therefore was likely a “cadenced rumble” ^16^. Three playback stimuli were elicited by another playback, and we assumed that the individual whose call was broadcast from the speaker was the intended receiver of the call that was produced in response to that playback.

We identified the receiver of natural calls as the only adult member of the family group who was separated from the caller during the call or the only individual who responded to the call. In one case, there were two adult females separated from the caller, and we assumed the receiver was the older of the two females who was in the lead and who rejoined the caller first (see Table S10). We prepared all playback stimuli in Audacity 3.0.2. Each stimulus consisted of a single rumble preceded by one second of background noise with a fade-in and followed by one second of background noise with a fade-out. In three cases, we applied a high-pass (5 Hz cutoff, 6 dB roll-off) or low-pass filter (1000 Hz cutoff, 6 dB roll-off) to remove excessive noise.

### Playback trial protocol

The stimuli were played back as .wav files (uncompressed audio) from an iPhone SE (Apple Inc., Cupertino, CA) attached to QLXD1 wireless bodypack transmitter (Shure, Niles, IL) transmitting to a custom-built loudspeaker (Bag End Loudspeakers, Algonguin, IL) (see Supplementary Information). We placed the speaker 40.2-59.0 m from the subject (mean 49.1 ± 4.2 m), either on the ground in front of a tree or shrub and covered by camouflage netting or on the edge of the rear seat of a Toyota double cab Landcruiser facing the door with all four doors and windows and both roof hatches open. Re-recordings at 50 m revealed no obvious difference between sounds played with the speaker on the ground or inside the vehicle. We only conducted playbacks when the original caller and “alternate receiver” (the other subject receiving playbacks from the same caller) were >180 m from and out of sight of the subject (>270 m from the alternate receiver if she had not yet received all her playbacks). When the original caller’s location was known (19/34 trials) the speaker was placed in approximately the same direction relative to the subject as the original caller. In the remaining trials the caller could not be located after searching a ∼300 m radius around the subject. Trials were redone after at least 7 days if the speaker malfunctioned, the subject moved her head out of sight right before the playback started, or we discovered after the playback that the speaker was not in the correct location relative to the subject and the original caller. During each trial we filmed the subject from inside the vehicle for at least 1 min, then played the stimulus once, and continued filming for at least another 10 min. We also recorded audio with an Earthworks QTC40 microphone and Sound Devices MixPre3-II recorder. The observers were blind to the playback condition (test or control) until all trials were complete and all videos and audio recordings were scored.

### Statistical analysis of playback data

From the video and audio recordings of each playback trial we measured the subject’s Latency to Approach the speaker, Latency to Vocalize, Number of Vocalizations produced within 10 min following the playback, Latency to Vigilance, and Change in Vigilance Duration in the minute following the playback compared to the minute preceding the playback. Latencies were defined as the time from the start of the playback until the behavior of interest occurred and were censored when the subject moved out of sight or at 10 min, whichever came first. Vigilance was defined as lifting head above shoulder level, moving head from side to side, holding ears away from body without flapping, or lifting trunk while sniffing toward speaker. We ran a separate model for each response variable with Subject ID as a random effect and Treatment and the following covariates/factors as fixed effects: Caller-Original Receiver Relationship (relationship between the caller and the original receiver of the call; Extended Data Table 3), Distance (distance in meters between the speaker and the subject), dBC (amplitude of the playback stimulus in dBC at 1 m), Other Adults (whether other adults were within 50 m of subject during playback), Speaker Location (whether speaker was on ground or in vehicle), and Cumulative Playback Exposure (cumulative number of playbacks to which subject was exposed at distance of 300 m or less, including trials that were redone and playbacks to other subjects). We used Cox proportional hazards regression in the coxme package ^38^ for the latency variables, a generalized linear model with a Poisson error distribution in the lme4 package ^39^ for Number of Vocalizations, and a linear model for Change in Vigilance Duration.

## ACKNOWLEDGEMENTS

We thank the Office of the President of Kenya, the Samburu, Isiolo, and Kajiado County governments, the Wildlife Research & Training Institute of Kenya, and Kenya Wildlife Service for permission to conduct fieldwork in Kenya. We thank Save The Elephants and the Amboseli Trust for Elephants for logistical support in the field, J. M. Leshudukule, D. M. Letitiya, and Norah Njiraini for assistance with the fieldwork, G. Pardo for blinding the playback stimuli, and S. Pardo for input on the statistical analyses. We thank J. Berger, W. Koenig, and A. Horn for comments on the manuscript. This project was funded by a Postdoctoral Research Fellowship in Biology to MP from the National Science Foundation (award no. 1907122) and grants to JP and PG from the National Geographic Society, Care for the Wild, and the Crystal Springs Foundation. Field work was supported by Save the Elephants.

## AUTHOR CONTRIBUTIONS

MP conceived the study. MP and DL collected the data in Samburu and JP and PG collected the data in Amboseli. MP and KF performed the statistical analysis and MP created the figures. MP drafted the manuscript and KF, JP, and GW edited it. CM, IDH, and GW provided resources and access to long-term datasets and GW supervised the study.

The authors declare no competing interests. Supplementary Information is available for this paper.

Correspondence and requests for materials should be addressed to MP. Reprints and permissions information is available at www.nature.com/reprints. Data are available at doi:10.5061/dryad.hmgqnk9nj Code is available at doi:10.5061/dryad.hmgqnk9nj

## TITLES AND LEGENDS FOR EXTENDED DATA

**Extended Data Figure 1. Violin plots illustrating distribution of data with respect to callers and receivers.** The dataset consisted of 625 total calls, 114 unique callers, and 119 unique receivers, but each caller only addressed a small number of the receivers in the dataset.

**Extended Data Figure 2. Schematic illustrating how spectral acoustic features were measured.** First, a spectrogram was calculated by applying a Fast Fourier Transform to the signal (Hamming window, 700 samples, 90% overlap). Then a mel filter bank with 26 overlapping triangular filters between 0-500 Hz was applied to each window of the spectrogram to produce a mel spectrogram. The mel spectrogram was then normalized by dividing the energy value in each cell by the total energy in that time window and these proportional energies were logit-transformed so they would not be limited to between 0 and 1. As features for the robust principal components analysis, we used the vector of energy in each of the 26 mel frequency bands as well as the vectors of delta and delta-delta values for each frequency band (representing the change and acceleration in energy over time, respectively). In the spectrogram and mel spectrogram in this figure, warmer colors indicate higher amplitudes (greater energy).

**Extended Figure 3. Scatterplots showing the separation in 3D space between calls from the same caller to different receivers.** Axes are the three most important variables for predicting receiver ID (means of PCs 33, 23, and 48) as determined from the variable importance scores of a conditional inference random forest using the spectral acoustic features. Each plot represents a single caller, each point is a single call, and receiver IDs are coded by both color and shape. This figure only includes calls where certainty of caller ID and receiver ID were at least 0.5 (no more than 2 possible candidates) and the caller made at least 3 calls each to at least 2 different receivers.

**Extended Data Table 1. Hypotheses and predictions tested in this study Extended Data Table 2. Acoustic features used in the random forest models**

All acoustic features were derived from either the sparse matrix or low-rank matrix of a robust principal components analysis performed on multiple acoustic contours of equal length that were measured directly from the signal. For the spectral acoustic features, the acoustic contours were the Hilbert amplitude envelope, the vector of energies in each of the 26 bands of a mel spectrogram, and the delta and delta-delta values of the mel spectral bands. For the cepstral acoustic features, the acoustic contours were the Hilbert amplitude envelope, first 12 mel-frequency cepstral coefficients, and the delta and delta-delta values of the first 12 cepstral coefficients. The principal components analysis was performed on a matrix of all the contours for each call stacked end-to-end.

**Extended Data Table 3. Definitions of social relationship categories between caller and receiver**

Categories were defined based on sex, age, and mother-offspring status, the most important factors influencing dominance and bond strength within an elephant family group. Females were defined as adults if ≥10 years old, and males were defined as adults if independent from their natal group. All non-adults under this definition were classified as juveniles. Six years was chosen as the cutoff for different age classes because it is between 1-2x the average inter-birth interval, so a female ≥6 years older than another individual could have been that individual’s allomother.

**Extended Data Table 4. Results for ANOVAs to test if calls with the same caller and receiver were more similar than calls with the same caller and different receivers**

Each observation was a pair of calls. ANOVA models were of the form Rank-transformed Proximity Score ∼ Same Caller Pair Type (whether the two calls in a pair had the same caller and receiver or same caller and different receivers) + Same Context (whether the two calls in a pair had the same behavioral context). Pairs of calls recorded on the same date or where the two calls had a different type of caller-receiver relationship were excluded. Three models were run for each set of acoustic features (spectral and cepstral): all pairs of calls meeting above criteria (n=2391), pairs of calls in which both calls were convergent on the receiver’s calls (n=252), and pairs of calls in which both calls were divergent from the receiver’s calls (n=798). Convergent calls = calls from caller A to receiver B that were more similar to receiver B’s calls than calls from caller A to other receivers were to receiver B’s calls. Divergent calls = calls from caller A to receiver B that were less similar to receiver B’s calls than calls from caller A to other receivers were to receiver B’s calls.

**Extended Data Table 5. Results for ANOVAs to test if calls addressed to a given receiver were imitative of the receiver’s calls**

Each observation was a pair of calls. ANOVA models were of the form Rank-transformed Proximity Score ∼ Imitation Pair Type + Same Relationship + Same Context + Caller Dyad ID. Model was run once for each set of acoustic features: spectral and cepstral. Imitation Pair Type = whether the caller of one call in a pair was the receiver of the other call. Same Relationship = whether the callers of both calls in a pair had the same type of relationship to their respective receivers. Same Context = whether the two calls in a pair were recorded in the same behavioral context (contact/greeting). Caller Dyad ID = identifier for the two callers in a pair. Pairs of calls recorded on the same date, from callers in different social groups, or with the same caller or receiver were excluded. We also excluded pairs of calls for which Caller Dyad ID only occurred with one level of Imitation Pair Type (final n=11,309).

**Extended Data Table 6. Results for ANOVAs to test if different callers used similar labels for the same receiver**

Each observation was a pair of calls. ANOVAs were of the form Rank-transformed Proximity Score ∼ Different Caller Pair Type + Same Relationship + Same Context. Model was run separately for each set of acoustic features: spectral and cepstral. Different Caller Pair Type = whether the two calls in a pair had different callers and the same receiver or different callers and different receivers. Same Relationship = whether the two calls in a pair had the same type of relationship between caller and receiver. Same Context = whether the two calls in a pair were recorded in the same behavioral context (contact/greeting) or not. Pairs of calls recorded on the same date or from callers in different social groups were excluded (final n=20,235)

**Extended Data Table 7. Summary of playback trials for each subject**

All callers and subjects were adult females except M25.0012 (subadult male) and M9.9612 (subadult female). The letter in parentheses after each caller ID represents a unique call (e.g., R23 (a) and R23 (b) were different calls recorded from R23). Twelve trials were redone once or twice because the playback system malfunctioned, the subject went out of sight just as the playback began, or the speaker was accidentally placed >60 m away or in the wrong direction relative to the subject and the original caller. Trials that were later redone are not included in this table.

