## Supplementary Information for "African elephants address one another with individually specific calls"

### SUPPLEMENTARY METHODS

#### Scoring behavioral context

For this study we only used rumbles produced in the contexts of "contact calling", "little-greeting", and "greeting", as these are the most common types of rumbles with a single receiver that are produced between adults [1]. We did not include rumbles from other behavioral contexts as these typically either involve multiple simultaneous receivers (e.g. "let's go" rumbles) or occur between calves and their caregiver (e.g. "begging" and "comforting" rumbles), allowing the receiver to be predicted simply from acoustic cues to caller identity and behavioral context, even if there were no vocal label identifying the intended receiver [1]. Nonetheless, there was a great deal of variation in the precise social context surrounding the production of each call and the age and internal state of the callers. As elephant rumbles vary with behavioral context, age, and the emotional state of the caller [1–3], this contextual heterogeneity of the recordings likely added substantial noise to the data.

We defined contact calling as calls produced by or addressed to an individual who was separated from the group and attempting to reinitiate contact. "Little-greeting" and "greeting" rumbles are both produced when one individual approaches another in an affiliative manner, but the latter are produced after a greater period of separation, are more likely to involve a face-to-face approach, and typically involve greater emotive behavior such as temporal gland streaming and pirouetting to stand in parallel [1]. We only used little-greeting rumbles from Amboseli as there were few non-overlapping greeting rumbles but included both contexts from Samburu as we did not clearly distinguish between greetings and little-greetings in Samburu. For the sake of brevity, all references to "greeting" rumbles in the main manuscript and from here on in the Supplemental Information refer collectively to both "greeting" and "little greeting" rumbles as defined by [1].

#### Scoring certainty of caller ID, behavioral context, and receiver ID

In Samburu, we recorded the certainty with which we knew caller ID, behavioral context, and receiver ID as 1 over the number of possible alternatives [4]. For example, in cases where we thought the call was plausibly addressed to a single individual but there were two possible candidates for who the receiver was, we designated one of the two individuals as the putative receiver and assigned the certainty of receiver ID a value of 0.5. In Amboseli, certainty of caller ID and behavioral context were originally scored as "certain", "fairly confident", "educated guess", "no idea". To facilitate comparison between the two datasets we rescored Amboseli values of "certain" or "fairly confident" as 1, "educated guess" as 0.5, and "no idea" as 0. When *ad lib* fieldnotes specified the number of possibilities for the caller ID or behavioral context of a call, we revised the certainty accordingly. Certainty of receiver ID was not systematically recorded in Amboseli. Unless the field notes specified that the receiver ID was uncertain, we assigned the certainty a value of 1. Behavioral observations were recorded by a single observer at each field site (MP in Samburu, JP in Amboseli). Since the observations at each field site were conducted in different years without accompanying video in most cases, there was no way to calculate inter-observer reliability.

#### Call segmentation

In Amboseli we wrote down the elapsed time on the recorder and contextual information for each call heard in the field; in Samburu we recorded verbal annotations onto a second channel of the recorder in real time using a Martel Stenomask, which isolated the sound of the observer's voice from the Earthworks microphone [4]. Using this information, we manually drew a selection box around the spectrogram of each call in Raven Pro 1.5 (Cornell Lab of Ornithology, Ithaca, NY), with a buffer of approximately 1 sec on either side of the call (Samburu (44.1 kHz sampling rate): Hann window, 50% overlap, window=11,878 samples, DFT=16,384 samples; Amboseli (2 kHz sampling rate): Hann window, 50% overlap, window=312 samples, DFT=512 samples). This automatically generated a selection table in .txt format with the filename and start and end times of each selection box, to which we added caller ID, receiver, ID, behavioral context, and the certainty of each. We performed all further acoustic and statistical analyses in R version 4.1.3 [5].

To determine the precise onset and offset of each call, we low-pass filtered the calls (Butterworth filter, order=5, cutoff=490 Hz), down-sampled them to 2000 Hz if not already at that sampling rate, applied a high-pass filter (Butterworth filter, order=10, cutoff=30 Hz), and normalized them to 70% of max amplitude and 16 bits of amplitude resolution using the packages seewave [6] and tuneR [7]. We then used the function segment() in the package soundgen [8] to detect the onset and offset of each call based on the amplitude envelope. We verified the automatically detected start and end time for each call by visual inspection of the amplitude envelope and spectrogram and manually adjusted the times when necessary.

**Acoustic measurements**

We trimmed the original unfiltered sound clips to the automatically detected start and end times, low-pass filtered the clips (Butterworth filter, order=5, cutoff=800 Hz), downsampled them to 2000 Hz if not already at that sampling rate, applied a high-pass filter (Butterworth filter, order=2, cutoff=4 Hz), and finally normalized them to 70% of the max amplitude and 16 bits of amplitude resolution. For each call, we measured the smoothed Hilbert amplitude envelope (moving average window, window length=350 ms, overlap=90%) and two alternative sets of features: normalized mel spectrogram and mel-frequency cepstral coefficients.

A mel spectrogram is similar to a traditional spectrogram (raster plot with time on the x-axis, frequency on the y-axis, and amplitude indicated by pixel darkness) but with frequency transformed to the logarithmic mel scale [9]. While the mel scale was designed to approximate human hearing sensitivity, most other mammals, including elephants, perceive frequency on a similar logarithmic scale [10]. We calculated a mel spectrogram for each call using the audspec() function of the tuneR package (26 mel-frequency bands between 0-500 Hz, 350 ms Hamming window, 90% overlap). We then normalized the mel spectrogram by dividing the energy value in each cell of the spectrogram by its column sum so that the energies would be a proportion of the total energy in each time window, and logit-transformed these proportional energies so the values would not be limited between 0 and 1. We also calculated delta and delta-delta values for each mel spectral band, with delta values being the differences between successive energy values in the mel spectral band (i.e., the change in energy over time within a mel spectral band), and delta-delta values being the differences between successive delta values (i.e., the acceleration of energy over time within a mel spectral band) (Fig. S2). We saved the vector of energies in each mel spectral band and their corresponding delta and delta-delta values as acoustic contours for further processing. While mel spectral bands have not previously been used as acoustic features for analyzing elephant calls, they describe more of the variation in the call than commonly used features such as fundamental frequency and formants, while remaining easily interpretable.

We also calculated mel-frequency cepstral coefficients (MFCCs) for each call, which are less interpretable than mel spectral bands but have been previously used successfully to classify elephant vocalizations [3,11,12]. MFCCs are calculated by applying a discrete cosine transform to each time window of a mel spectrogram, with the coefficients of the discrete cosine transform being the cepstral coefficients [13]. Each cepstral coefficient can be thought of as representing the degree of modulation of the spectrum at a different period, with lower numbered coefficients representing slower periods of modulation. Since MFCCs are calculated for each time window of the mel spectrogram, the output is a vector of values for each cepstral coefficient. We calculated MFCCs using the melfcc() function in the tuneR package, with a time window of 350 ms with 90% overlap, 40 mel-frequency bands between 0-500 Hz, and a pre-emphasis filter with a cutoff frequency of 10 Hz, and kept the first 12 coefficients (12 vectors per call) for further processing. We also calculated delta and delta-delta values for the first 12 cepstral coefficient contours.

**Extraction of derived features from acoustic contours**

We rescaled the acoustic contours by arranging them in a matrix with each contour in a separate row, and then subtracting the column median from each value and dividing the result by the column mean average deviation. We decorrelated the contours with robust principal components analysis in the rpca package in R, which separates the data into a low-rank matrix of robust principal components without outliers, and a sparse matrix containing the outlier values ($\lambda$=0.00996) [14]. Robust PCA has the advantage over standard PCA of being more resilient to noisy data. We extracted four measurements from the sparse matrix to use for statistical analysis: median, robust skewness, and two measures of spread: minimum extent and equivalent statistical extent. We also calculated the means of the first *n* low-rank principal components required to explain 99.9% of the variation (74 for spectral features, 12 for cepstral features).

We used multi-taper spectral estimation [15] to derive the frequency spectra of the low-rank principal components that explained 99.9% of the variation (treating each principal component as if it were a waveform) and calculated an F-ratio for each point in each spectrum, testing the null hypothesis that the spectral value in question could have been derived from a random waveform. We calculated the mean of the F-ratios at each point across the aligned spectra and selected the four largest peaks in the series of mean F-ratios. We sorted these peaks in order of increasing frequency and calculated the frequency and magnitude of each peak.

We calculated the same metrics on spectra that were weighted according to the proportion of variation that was explained by the principal component from which the spectrum was derived. We multiplied the F-ratios in each of the spectra by the proportion of variation in the data explained by the principal component in question, summed the weighted F-ratios at each point in the aligned spectra, and then calculated the frequencies and magnitudes of the four largest peaks in the summed F-ratios, sorted in order of increasing frequency. The final acoustic features used in our models are summarized in Table S2. We ran all subsequent statistical analyses separately for the spectral and cepstral acoustic features.

**Checking ANOVA model assumptions**

For all rank-transformed ANOVA models, we checked the assumption of normality by visually examining Q-Q plots and histograms of the residuals. We checked the assumption of equal variances by visually examining a boxplot of all groups. The residuals for all models deviated from normality with relatively flat-topped distributions (Fig. S5). However, ANOVA has been shown to be robust even to severe deviations from normality with skewness as high as 2 and excess kurtosis as high as 6 (a normal distribution has a skewness of 0 and excess kurtosis of 0) [16]. Skewness of the residuals in our models was between -0.031 and 0.066 and excess kurtosis was between -1.19 and -0.98, so we deemed ANOVA an appropriate choice of model despite the deviations from normality. Boxplots indicated approximately equal variances for all groups (Fig. S6).

#### Playback system and volume

The cord connecting the playback device to the wireless transmitter had to be replaced three times over the course of the experiment, each time changing the output level of the speaker. Thus, depending on which cord was in use, we normalized the stimuli to -24, -22.5, or -18 dB in Audacity 3.0.2 to ensure a functionally equivalent normalization level across all trials.

The speaker's frequency response was flat from 10 Hz to 500 Hz up to a given maximum output level (maximum output = 89 dB SPL at 10 Hz, 101 dB SPL at 20 Hz, 113 dB SPL at 40 Hz). If the signal exceeded the maximum output at a given frequency, the speaker automatically reduced the level of the frequencies in question to avoid damage. Reported amplitudes for natural contact calls range from 94 to 115 dB SPL (extrapolated value at 1 m from source) [1,17]. We did not have access to an SPL meter with a flat frequency response at low frequencies, but our playback stimuli ranged from 96.2 to 104.3 dBC at 1m measured with a Protmex PT6708 sound level meter (Protech International Group Co., Shenzhen, China) or 93.4 to 102.9 dB SPL at 1m measured with the SoundMeter 10.5.8 iPhone application (Faber Acoustical, Lehi, UT). Mean measured volume did not differ between test and control stimuli (dBC: t-test, $t_{32.0}=0.03$, $p=0.97$; dB SPL: t-test, $t_{32.0}=0.15$, $p=0.88$).

### SUPPLEMENTARY DISCUSSION

#### Variable importance scores

Many of the same acoustic features scored highly in importance for predicting both caller and receiver ID, although the exact order of importance differed between the two models (Tables S1, S2). This suggests that rather than being encoded in entirely separate acoustic features, caller ID and receiver ID may both be conveyed by an interaction among many of the same acoustic parameters. However, given the partial aliasing of caller ID and receiver ID in our dataset (Fig. S1), it is also possible that the model sometimes used features associated with caller ID to predict receiver ID (especially for calls that lacked a vocal label of receiver ID), which could make the top features for the two models more similar than they would be otherwise.

The top features for predicting caller and receiver ID were derived from principal components that explained a small fraction of variance in the data. The top feature for both receiver ID and caller ID was the mean of the 33$^{rd}$ principal component, which explained only 0.78% of the variance in the data, and the top 5/94 features for both models were means of principal components that collectively explained only 3.59% (caller ID) and 3.36% (receiver ID) of the variance (Table S1). This suggests that cues to caller and receiver ID account for only a small proportion of the total variation in rumbles, which likely explains why they cannot be easily ascertained by visual inspection of the spectrogram. Rumbles also convey information on behavioral context and the internal state of the caller [1,18], and these factors may account for a larger proportion of the variation in rumbles than signals of caller or receiver ID.

To interpret the variable importance scores for the principal components in terms of the acoustic properties of the call, we related the principal components back to the original acoustic contours used to calculate them (mel frequency bands and their delta and delta-delta values, and amplitude envelope). Several of the mel frequency bands represented in the top ten acoustic contours for predicting caller ID, receiver ID, or both were in the ranges of 20-50 Hz and 60-200 Hz, which correspond to the locations of the first and second formants, respectively, in most rumbles produced by adult African savannah elephants [19,20] (Table S2). Previous studies have found the first and second formant of elephant rumbles to be important for encoding a variety of information, including caller identity, social context, and the presence of external threats [2,19,21]. While the top acoustic contours associated with the first formant in our study were the mel frequency bands themselves (i.e., the amount of energy in that frequency range), the top acoustic contours associated with the second formant were delta-delta coefficients (i.e., the rapidity of change in energy within that frequency range). Most studies that have measured formants in elephant rumbles measured a single value for each formant, either by measuring over a short interval in the middle of the call [2,18,20,21] or averaging across the whole call [19]. Our results suggest that modulation of the formants over time is likely important for encoding certain types of information and should be measured as well.

Six of the top ten acoustic contours for predicting receiver ID and four of the top ten for predicting caller ID were delta-delta coefficients (acceleration of energy) for mel frequency bands above 300 Hz (Table S9). This was unexpected, as many elephant rumbles have little energy in this frequency range and several prior studies have ignored energy above 300 Hz altogether [2,18,21,22]. It may be that higher frequencies were particularly informative for discriminating among calls in our random forest model precisely because only some calls have appreciable energy in this bandwidth. Nonetheless, this result suggests that ignoring higher frequencies in acoustic analysis of elephant rumbles may risk missing important information.

**Table S1. Variable importance scores of spectral acoustic features, calculated from a conditional inference random forest model**

| Receiver ID | | | Caller ID | | |
|---|---|---|---|---|---|
| ***Feature*** | ***Prop. of Variance*** | ***Importance*** | ***Feature*** | ***Prop. of Variance*** | ***Importance*** |
| Mean PC33 | 0.0077605 | 0.000372 | Mean PC33 | 0.0077605 | 0.000596 |
| Mean PC23 | 0.0111941 | 0.000204 | Mean PC49 | 0.0049136 | 0.000304 |
| Mean PC48 | 0.0049651 | 0.000188 | Mean PC23 | 0.0111941 | 0.000288 |
| Mean PC37 | 0.0070876 | 0.00018 | Mean PC37 | 0.0070876 | 0.000284 |
| Mean PC61 | 0.002563 | 0.000176 | Mean PC48 | 0.0049651 | 0.000272 |
| Mean PC22 | 0.0118438 | 0.000172 | Mean PC57 | 0.0030659 | 0.000256 |
| Mean PC1 | 0.1585693 | 0.00016 | Mean PC17 | 0.0145093 | 0.000252 |
| Mean PC62 | 0.0024693 | 0.000144 | Mean PC1 | 0.1585693 | 0.00022 |
| Mean PC34 | 0.0076116 | 0.000108 | Mean PC61 | 0.002563 | 0.000172 |
| Mean PC72 | 0.0007939 | 0.000108 | Mean PC62 | 0.0024693 | 0.000156 |
| Mean PC64 | 0.0022598 | 0.000104 | Mean PC66 | 0.002015 | 0.00014 |
| Mean PC60 | 0.0026775 | 9.2e-05 | Mean PC24 | 0.0110066 | 0.000124 |
| Mean PC24 | 0.0110066 | 8.8e-05 | Mean PC34 | 0.0076116 | 0.00012 |
| Mean PC49 | 0.0049136 | 7.6e-05 | Mean PC22 | 0.0118438 | 0.000116 |
| Mean PC74 | 0.0005722 | 7.6e-05 | Mean PC43 | 0.0060188 | 0.000108 |
| Mean PC38 | 0.00683 | 7.2e-05 | Mean PC41 | 0.0063804 | 0.000104 |
| Mean PC42 | 0.0061708 | 6e-05 | Mean PC3 | 0.0811011 | 0.000104 |
| Mean PC2 | 0.0946066 | 5.6e-05 | Mean PC32 | 0.0080467 | 9.6e-05 |
| Mean PC26 | 0.0102873 | 5.6e-05 | Mean PC16 | 0.0152261 | 8.4e-05 |
| Rob. skew. (sparse) | NA | 5.6e-05 | Mean PC20 | 0.0129081 | 8.4e-05 |
| Mean PC66 | 0.002015 | 5.6e-05 | Mean PC29 | 0.0089573 | 8e-05 |
| Mag. peak4 | NA | 5.6e-05 | Mean PC28 | 0.009652 | 7.6e-05 |
| Mean PC57 | 0.0030659 | 5.2e-05 | Mean PC26 | 0.0102873 | 7.6e-05 |
| Mean PC71 | 0.0009098 | 4.8e-05 | Mean PC68 | 0.0017909 | 7.2e-05 |
| Mean PC51 | 0.0042509 | 4.8e-05 | Mean PC45 | 0.0054391 | 7.2e-05 |
| Mean PC20 | 0.0129081 | 4.4e-05 | Mean PC69 | 0.001351 | 7.2e-05 |
| Mean PC30 | 0.0087972 | 4.4e-05 | Mean PC7 | 0.0302463 | 7.2e-05 |
| Mean PC69 | 0.001351 | 4.4e-05 | Mean PC74 | 0.0005722 | 7.2e-05 |
| Mean PC10 | 0.0216232 | 4e-05 | Mean PC60 | 0.0026775 | 6.8e-05 |
| Mean PC53 | 0.0039579 | 4e-05 | Mean PC19 | 0.0130311 | 6.8e-05 |
| Mean PC14 | 0.0169985 | 3.6e-05 | Mean PC40 | 0.00663 | 6.8e-05 |
| Mean PC18 | 0.0132465 | 3.6e-05 | Mean PC64 | 0.0022598 | 6.8e-05 |
| Mean PC52 | 0.0041949 | 3.6e-05 | Mean PC15 | 0.0153567 | 6.4e-05 |
| Mean PC15 | 0.0153567 | 3.2e-05 | Mag. peak1 (weighted) | NA | 6e-05 |
| Mean PC31 | 0.0086248 | 3.2e-05 | Mag. peak2 (weighted) | NA | 6e-05 |

| | | | | | |
|---|---|---|---|---|---|
| Mag. peak4 (weighted) | NA | 3.2e-05 | Mean PC9 | 0.0224639 | 6e-05 |
| Min. ext. (sparse) | NA | 3.2e-05 | Mean PC54 | 0.003751 | 6e-05 |
| Mean PC25 | 0.0106067 | 2.8e-05 | Mean PC53 | 0.0039579 | 6e-05 |
| Mean PC44 | 0.0057159 | 2.8e-05 | Mean PC31 | 0.0086248 | 5.6e-05 |
| Mean PC6 | 0.0329591 | 2.8e-05 | Mean PC30 | 0.0087972 | 5.2e-05 |
| Mean PC3 | 0.0811011 | 2.8e-05 | Mean PC55 | 0.0035533 | 4.8e-05 |
| Mean PC7 | 0.0302463 | 2.8e-05 | Mean PC13 | 0.017768 | 4.8e-05 |
| Mean PC8 | 0.0249251 | 2.8e-05 | Mean PC14 | 0.0169985 | 4.4e-05 |
| Mean PC43 | 0.0060188 | 2.8e-05 | Mean PC18 | 0.0132465 | 4.4e-05 |
| Mag. peak3 | NA | 2.8e-05 | Freq. peak2 (weighted) | NA | 4e-05 |
| Mean PC29 | 0.0089573 | 2.4e-05 | Mean PC10 | 0.0216232 | 4e-05 |
| Mean PC28 | 0.009652 | 2.4e-05 | Mean PC11 | 0.0212398 | 4e-05 |
| Mean PC4 | 0.0434407 | 2.4e-05 | Median (sparse) | NA | 4e-05 |
| Mean PC19 | 0.0130311 | 2.4e-05 | Mag. peak3 (weighted) | NA | 3.6e-05 |
| Mag. peak1 (weighted) | NA | 2.4e-05 | Rob. skew. (sparse) | NA | 3.6e-05 |
| Mean PC9 | 0.0224639 | 2.4e-05 | Mean PC39 | 0.0066973 | 3.6e-05 |
| Mean PC11 | 0.0212398 | 2e-05 | Mean PC44 | 0.0057159 | 3.2e-05 |
| Mean PC17 | 0.0145093 | 2e-05 | Mean PC2 | 0.0946066 | 2.8e-05 |
| Mean PC40 | 0.00663 | 2e-05 | Mean PC38 | 0.00683 | 2.8e-05 |
| Mean PC27 | 0.0098732 | 2e-05 | Mean PC6 | 0.0329591 | 2.8e-05 |
| Mean PC35 | 0.0074461 | 2e-05 | Mean PC4 | 0.0434407 | 2.4e-05 |
| Mean PC41 | 0.0063804 | 1.6e-05 | Mean PC21 | 0.0123849 | 2.4e-05 |
| Mean PC67 | 0.0019018 | 1.6e-05 | Mean PC70 | 0.0012292 | 2.4e-05 |
| Mean PC13 | 0.017768 | 1.6e-05 | Freq. peak2 | NA | 2.4e-05 |
| Freq. peak3 | NA | 1.2e-05 | Mean PC56 | 0.0033799 | 2e-05 |
| Mean PC47 | 0.0051982 | 1.2e-05 | Mean PC12 | 0.0199427 | 2e-05 |
| Mean PC12 | 0.0199427 | 1.2e-05 | Mag. peak1 | NA | 2e-05 |
| Mean PC63 | 0.0023531 | 1.2e-05 | Mean PC8 | 0.0249251 | 2e-05 |
| Mean PC73 | 0.0006832 | 1.2e-05 | Mean PC52 | 0.0041949 | 2e-05 |
| Mean PC54 | 0.003751 | 1.2e-05 | Mean PC73 | 0.0006832 | 1.6e-05 |
| Median (sparse) | NA | 8e-06 | Mag. peak4 (weighted) | NA | 1.6e-05 |
| Mean PC50 | 0.004473 | 8e-06 | Mean PC46 | 0.005303 | 1.6e-05 |
| Mag. peak3 (weighted) | NA | 8e-06 | Mean PC47 | 0.0051982 | 1.6e-05 |
| Freq. PC peak4 | NA | 8e-06 | Mean PC36 | 0.007161 | 1.6e-05 |
| Mean PC32 | 0.0080467 | 8e-06 | Mean PC71 | 0.0009098 | 1.6e-05 |
| Freq. peak1 (weighted) | NA | 8e-06 | Mean PC72 | 0.0007939 | 1.2e-05 |

| | | | | | |
|---|---|---|---|---|---|
| Freq. PC peak2 | NA | 8e-06 | Mean PC42 | 0.0061708 | 1.2e-05 |
| Eq. stat. ext. (sparse) | NA | 4e-06 | Freq. peak3 (weighted) | NA | 1.2e-05 |
| Mean PC45 | 0.0054391 | 0 | Min. ext. (sparse) | NA | 1.2e-05 |
| Mean PC21 | 0.0123849 | 0 | Mag. peak4 | NA | 8e-06 |
| Mean PC56 | 0.0033799 | 0 | Mean PC63 | 0.0023531 | 8e-06 |
| Mag. PC peak2 | NA | 0 | Mean PC65 | 0.0022536 | 8e-06 |
| Mean PC36 | 0.007161 | -4e-06 | Mean PC5 | 0.0374864 | 8e-06 |
| Mean PC5 | 0.0374864 | -4e-06 | Mean PC67 | 0.0019018 | 4e-06 |
| Mean PC55 | 0.0035533 | -8e-06 | Freq. peak4 | NA | 4e-06 |
| Mag. peak2 (weighted) | NA | -8e-06 | Freq. peak1 | NA | 0 |
| Mag. peak1 | NA | -1.2e-05 | Freq. peak4 (weighted) | NA | -4e-06 |
| Mean PC46 | 0.005303 | -1.6e-05 | Mean PC51 | 0.0042509 | -4e-06 |
| Mean PC65 | 0.0022536 | -1.6e-05 | Mean PC35 | 0.0074461 | -8e-06 |
| Mean PC39 | 0.0066973 | -1.6e-05 | Mean PC50 | 0.004473 | -8e-06 |
| Freq. peak4 (weighted) | NA | -1.6e-05 | Mean PC25 | 0.0106067 | -8e-06 |
| Mean PC58 | 0.003017 | -2e-05 | Mean PC27 | 0.0098732 | -1.2e-05 |
| Mean PC68 | 0.0017909 | -3.2e-05 | Eq. stat. ext. (sparse) | NA | -1.2e-05 |
| Freq. peak3 (weighted) | NA | -4e-05 | Mag. peak2 | NA | -2.4e-05 |
| Freq. peak1 | NA | -4e-05 | Mag. peak3 | NA | -2.4e-05 |
| Mean PC16 | 0.0152261 | -4.4e-05 | Mean PC59 | 0.0029531 | -3.2e-05 |
| Freq. peak2 (weighted) | NA | -4.8e-05 | Freq. peak3 | NA | -3.6e-05 |
| Mean PC59 | 0.0029531 | -4.8e-05 | Freq. peak1 (weighted) | NA | -4e-05 |
| Mean PC70 | 0.0012292 | -6.8e-05 | Mean PC58 | 0.003017 | -7.6e-05 |

Variable importance scores calculated separately for model predicting receiver ID and model predicting caller ID. For features that were means of principal components (PCs), the proportion of variance in the data explained by the PC in question is given. Features were derived by stacking the matrix of acoustic contours (mel spectral bands, delta and delta-delta values, and amplitude envelope) for each call end to end and decomposing this matrix into a low-rank and sparse matrix using robust PCA. Features labeled "Mean PC" indicate the mean for a particular low-rank PC. Features labeled "Mag. peak" or "Freq. peak" indicate the magnitude or frequency, respectively, of one of the four largest spectral peaks (1=largest, 4=smallest) in the low-rank PCs, calculated by multi-taper spectral estimation on the aligned PCs. Magnitudes and frequencies labeled "(weighted)" were calculated from PC spectra weighted by the proportion of variation explained by each PC. Features labeled “(sparse)” were calculated from the sparse matrix (median, robust skewness, minimum extent, and equivalent statistical extent).

**Table S2. Ordered importance (most important to least) of the original spectral acoustic contours inferred from the loading of each acoustic feature on each low-rank principal component and the variable importance score of the mean of the principal component in question**

| Order of importance | Receiver ID | | | Caller ID | | |
|---|---|---|---|---|---|---|
| | *Acoustic feature* | *Frequency range (Hz)* | *Weighted sum* | *Acoustic feature* | *Frequency range (Hz)* | *Weighted sum* |
| 1 | DD22 | 365-408 | 0.0004280 | DD6 | 73-105 | 0.0006520 |
| 2 | MelBand2 | 14-43 | 0.0004186 | DD24 | 408-453 | 0.0006399 |
| 3 | DD19 | 303-344 | 0.0004078 | DD22 | 365-408 | 0.0006129 |
| 4 | DD24 | 408-453 | 0.0004066 | DD10 | 138-172 | 0.0006076 |
| 5 | DD26 | 453-500 | 0.0003929 | DD12 | 172-207 | 0.0005963 |
| 6 | DD21 | 344-386 | 0.0003866 | DD26 | 453-500 | 0.0005903 |
| 7 | DD6 | 73-105 | 0.0003819 | DD11 | 155-189 | 0.0005858 |
| 8 | DD12 | 172-207 | 0.0003784 | DD19 | 303-344 | 0.0005857 |
| 9 | DD23 | 386-430 | 0.0003751 | MelBand2 | 14-43 | 0.0005829 |
| 10 | MelBand3 | 29-58 | 0.0003736 | DD15 | 226-263 | 0.0005827 |
| 11 | MelBand5 | 58-89 | 0.0003715 | DD21 | 344-386 | 0.0005811 |
| 12 | DD15 | 226-263 | 0.0003711 | DD7 | 89-121 | 0.0005658 |
| 13 | DD17 | 263-303 | 0.0003709 | MelBand5 | 58-89 | 0.0005621 |
| 14 | DD11 | 155-189 | 0.0003685 | MelBand3 | 29-58 | 0.0005611 |
| 15 | MelBand12 | 172-207 | 0.0003678 | Delta26 | 453-500 | 0.0005585 |
| 16 | MelBand14 | 207-244 | 0.0003645 | DD23 | 386-430 | 0.0005541 |
| 17 | DD10 | 138-172 | 0.0003633 | MelBand13 | 189-226 | 0.0005524 |
| 18 | MelBand9 | 121-155 | 0.0003620 | DD17 | 263-303 | 0.0005511 |
| 19 | DD14 | 207-244 | 0.0003601 | MelBand4 | 43-73 | 0.0005477 |
| 20 | MelBand17 | 263-303 | 0.0003600 | MelBand26 | 453-500 | 0.0005427 |
| 21 | MelBand7 | 89-121 | 0.0003593 | Delta21 | 344-386 | 0.0005363 |
| 22 | MelBand10 | 138-172 | 0.0003559 | MelBand9 | 121-155 | 0.0005315 |
| 23 | MelBand26 | 453-500 | 0.0003547 | Delta11 | 155-189 | 0.0005276 |
| 24 | DD18 | 283-323 | 0.0003544 | MelBand12 | 172-207 | 0.0005265 |
| 25 | DD20 | 323-365 | 0.0003394 | DD3 | 29-58 | 0.0005242 |
| 26 | MelBand13 | 189-226 | 0.0003360 | Delta17 | 263-303 | 0.0005237 |
| 27 | DD9 | 121-155 | 0.0003357 | DD13 | 189-226 | 0.0005186 |
| 28 | DD3 | 29-58 | 0.0003321 | DD14 | 207-244 | 0.0005106 |
| 29 | MelBand4 | 43-73 | 0.0003321 | DD18 | 283-323 | 0.0005093 |
| 30 | DD5 | 58-89 | 0.0003287 | DD9 | 121-155 | 0.0005081 |
| 31 | MelBand8 | 105-138 | 0.0003242 | Delta9 | 121-155 | 0.0004998 |
| 32 | MelBand6 | 73-105 | 0.0003231 | Delta13 | 189-226 | 0.0004987 |
| 33 | Delta21 | 344-386 | 0.0003180 | MelBand14 | 207-244 | 0.0004979 |
| 34 | DD7 | 89-121 | 0.0003175 | MelBand8 | 105-138 | 0.0004908 |
| 35 | DD13 | 189-226 | 0.0003168 | Delta12 | 172-207 | 0.0004872 |
| 36 | Delta26 | 453-500 | 0.0003132 | MelBand10 | 138-172 | 0.0004866 |
| 37 | MelBand20 | 323-365 | 0.0003132 | DD20 | 323-365 | 0.0004860 |

| | | | | | | |
|---|---|---|---|---|---|---|
| 38 | Delta13 | 189-226 | 0.0003099 | Delta24 | 408-453 | 0.0004807 |
| 39 | MelBand19 | 303-344 | 0.0003066 | Delta15 | 226-263 | 0.0004791 |
| 40 | Delta17 | 263-303 | 0.0003027 | DD5 | 58-89 | 0.0004785 |
| 41 | DD16 | 244-283 | 0.0003016 | MelBand15 | 226-263 | 0.0004725 |
| 42 | MelBand23 | 386-430 | 0.0003000 | Delta20 | 323-365 | 0.0004723 |
| 43 | Delta9 | 121-155 | 0.0002977 | Delta23 | 386-430 | 0.0004722 |
| 44 | DD25 | 430-476 | 0.0002959 | Delta10 | 138-172 | 0.0004721 |
| 45 | MelBand24 | 408-453 | 0.0002946 | MelBand6 | 73-105 | 0.0004706 |
| 46 | MelBand21 | 344-386 | 0.0002933 | DD16 | 244-283 | 0.0004704 |
| 47 | Delta7 | 89-121 | 0.0002904 | MelBand17 | 263-303 | 0.0004656 |
| 48 | MelBand15 | 226-263 | 0.0002903 | Delta16 | 244-283 | 0.0004589 |
| 49 | Delta12 | 172-207 | 0.0002901 | DD4 | 43-73 | 0.0004587 |
| 50 | Delta20 | 323-365 | 0.0002900 | MelBand7 | 89-121 | 0.0004554 |
| 51 | MelBand22 | 365-408 | 0.0002890 | Delta22 | 365-408 | 0.0004480 |
| 52 | Delta18 | 283-323 | 0.0002877 | Delta14 | 207-244 | 0.0004434 |
| 53 | MelBand25 | 430-476 | 0.0002863 | MelBand16 | 244-283 | 0.0004407 |
| 54 | Delta15 | 226-263 | 0.0002825 | Delta7 | 89-121 | 0.0004376 |
| 55 | MelBand1 | 0-29 | 0.0002820 | MelBand25 | 430-476 | 0.0004367 |
| 56 | Delta11 | 155-189 | 0.0002804 | DD8 | 105-138 | 0.0004360 |
| 57 | Delta16 | 244-283 | 0.0002784 | DD25 | 430-476 | 0.0004324 |
| 58 | MelBand16 | 244-283 | 0.0002742 | MelBand24 | 408-453 | 0.0004250 |
| 59 | Delta10 | 138-172 | 0.0002733 | Delta18 | 283-323 | 0.0004230 |
| 60 | MelBand18 | 283-323 | 0.0002703 | Delta25 | 430-476 | 0.0004179 |
| 61 | Delta23 | 386-430 | 0.0002623 | MelBand1 | 0-29 | 0.0004175 |
| 62 | Delta24 | 408-453 | 0.0002612 | Delta19 | 303-344 | 0.0004028 |
| 63 | MelBand11 | 155-189 | 0.0002595 | MelBand23 | 386-430 | 0.0004004 |
| 64 | Delta14 | 207-244 | 0.0002595 | Delta8 | 105-138 | 0.0004002 |
| 65 | DD8 | 105-138 | 0.0002555 | MelBand22 | 365-408 | 0.0003981 |
| 66 | Delta25 | 430-476 | 0.0002550 | Delta4 | 43-73 | 0.0003960 |
| 67 | Delta4 | 43-73 | 0.0002482 | MelBand19 | 303-344 | 0.0003943 |
| 68 | Delta22 | 365-408 | 0.0002481 | MelBand20 | 323-365 | 0.0003936 |
| 69 | DD2 | 14-43 | 0.0002480 | Delta5 | 58-89 | 0.0003935 |
| 70 | DD4 | 43-73 | 0.0002363 | MelBand21 | 344-386 | 0.0003902 |
| 71 | Delta5 | 58-89 | 0.0002338 | Delta6 | 73-105 | 0.0003675 |
| 72 | Delta3 | 29-58 | 0.0002178 | Delta3 | 29-58 | 0.0003472 |
| 73 | Delta6 | 73-105 | 0.0002160 | DD2 | 14-43 | 0.0003465 |
| 74 | Delta19 | 303-344 | 0.0002160 | MelBand18 | 283-323 | 0.0003342 |
| 75 | Delta8 | 105-138 | 0.0002023 | MelBand11 | 155-189 | 0.0003267 |
| 76 | AmpEnv | NA | 0.0001861 | AmpEnv | NA | 0.0003150 |
| 77 | DD1 | 0-29 | 0.0001785 | DD1 | 0-29 | 0.0002847 |
| 78 | Delta2 | 14-43 | 0.0001661 | Delta2 | 14-43 | 0.0002436 |
| 79 | Delta1 | 0-29 | 0.0001289 | Delta1 | 0-29 | 0.0001691 |

Each acoustic feature is a vector of values. Features labeled “MelBand” are the energy in each time frame of a given mel frequency band relative to the total energy in that time frame, features labeled “Delta” are the difference between energies in successive time frames of a mel frequency band (i.e., the change in energy over time), features labeled “DD” (delta-delta) are the difference between successive delta values for a mel frequency band (i.e., the acceleration of energy), and “AmpEnv” is the Hilbert amplitude envelope. There are 26 mel frequency bands, numbered from lowest to highest frequency, which increase in bandwidth with frequency. Frequency range (Hz) indicates the frequencies in Hertz spanned by a particular mel frequency band. Weighted sum was calculated by multiplying the variable’s loading on each principal component by the variable importance score of the mean of the principal component in question, and then summing up the weighted loadings for each variable (higher weighted sum = greater importance.
