## Extended Figures and Tables for "African elephants address one another with individually specific calls"

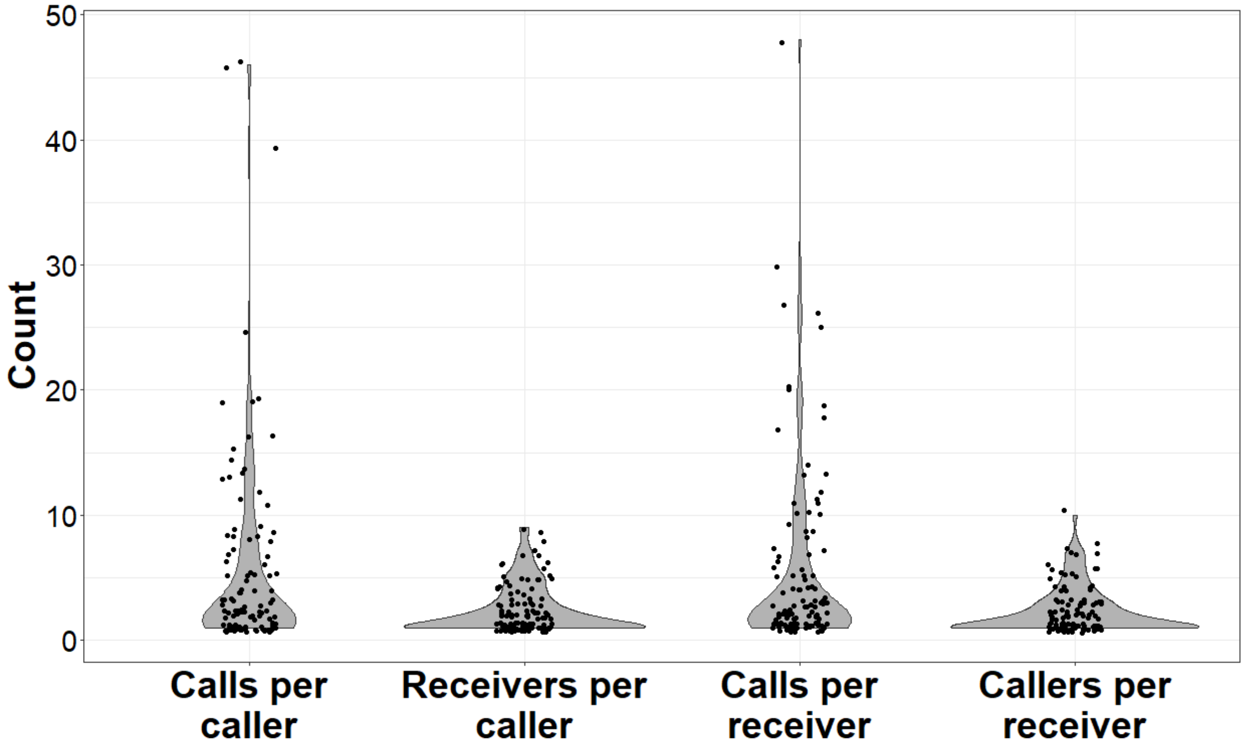

**Extended Data Figure 1. Violin plots illustrating distribution of data with respect to callers and receivers.** The dataset consisted of 625 total calls, 114 unique callers, and 119 unique receivers, but each caller only addressed a small number of the receivers in the dataset.

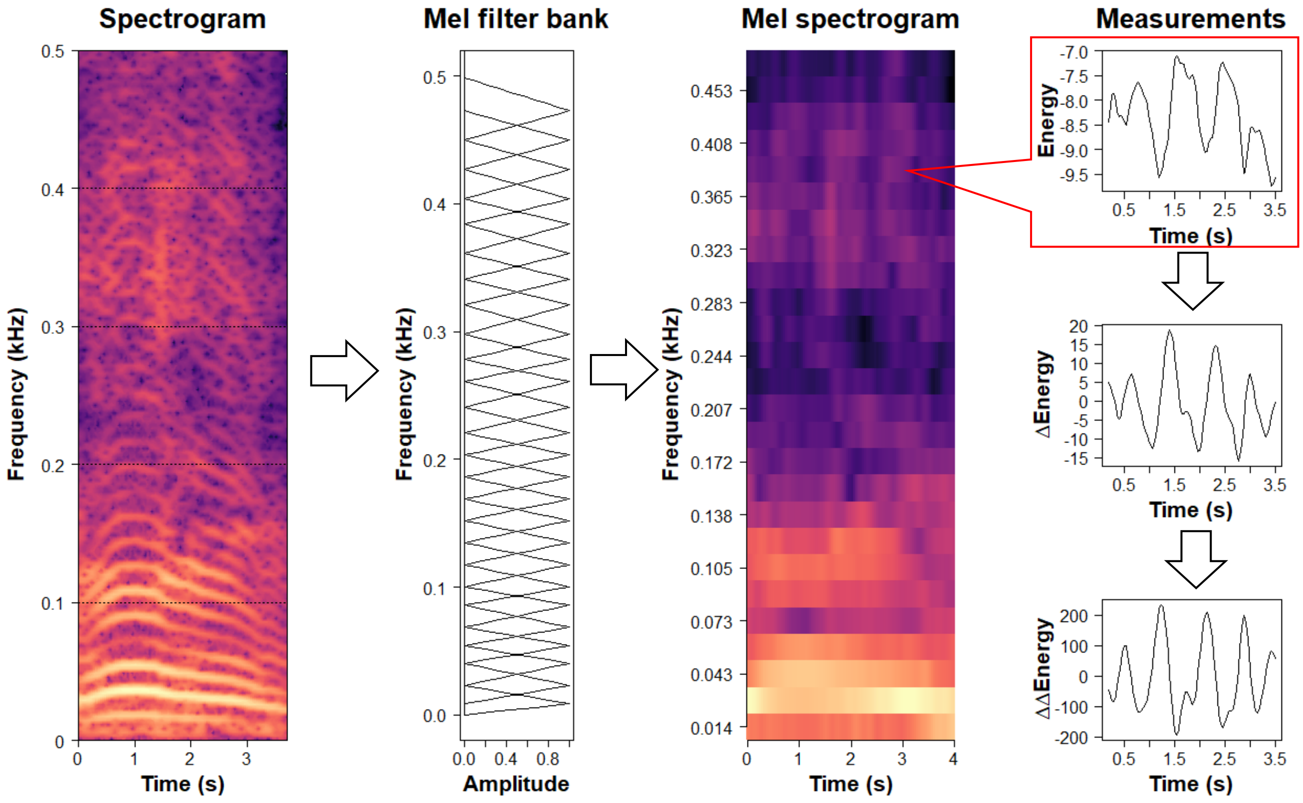

**Extended Data Figure 2. Schematic illustrating how spectral acoustic features were measured.** First, a spectrogram was calculated by applying a Fast Fourier Transform to the signal (Hamming window, 700 samples, 90% overlap). Then a mel filter bank with 26 overlapping triangular filters between 0-500 Hz was applied to each window of the spectrogram to produce a mel spectrogram. The mel spectrogram was then normalized by dividing the energy value in each cell by the total energy in that time window and these proportional energies were logit-transformed so they would not be limited to between 0 and 1. As features for the robust principal components analysis, we used the vector of energy in each of the 26 mel frequency bands as well as the vectors of delta and delta-delta values for each frequency band (representing the change and acceleration in energy over time, respectively). In the spectrogram and mel spectrogram in this figure, warmer colors indicate higher amplitudes (greater energy).

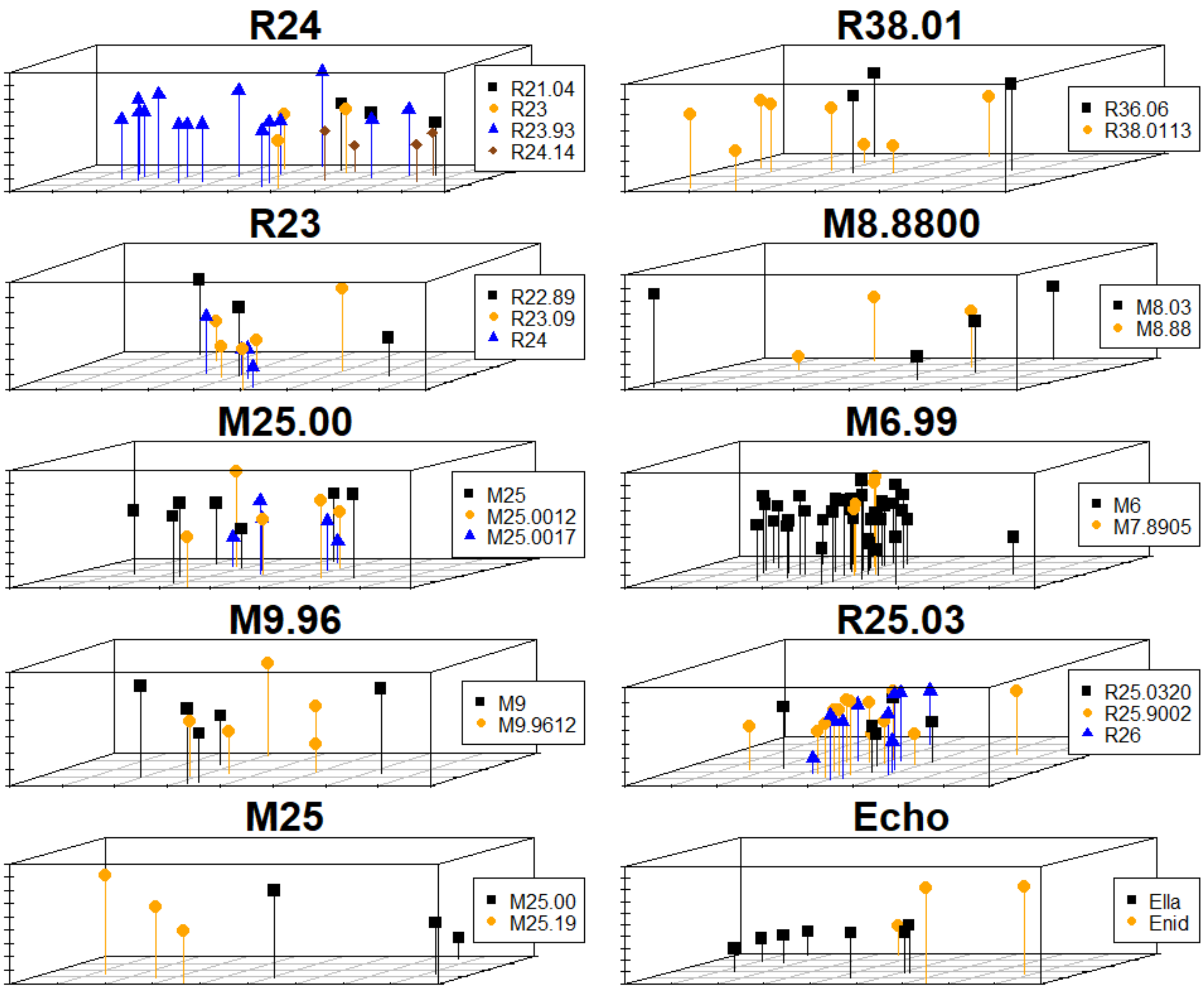

**Extended Figure 3. Scatterplots showing the separation in 3D space between calls from the same caller to different receivers.** Axes are the three most important variables for predicting receiver ID (means of PCs 33, 23, and 48) as determined from the variable importance scores of a conditional inference random forest using the spectral acoustic features. Each plot represents a single caller, each point is a single call, and receiver IDs are coded by both color and shape. This figure only includes calls where certainty of caller ID and receiver ID were at least 0.5 (no more than 2 possible candidates) and the caller made at least 3 calls each to at least 2 different receivers.

**Extended Table 1. Hypotheses and predictions tested in this study**

| Hypotheses | Predictions |
|---|---|
| 1. Elephants vocally label individual conspecifics | 1. Receiver ID can be predicted from call structure<br>2. Calls with same caller and same receiver will be more similar than calls with same caller and different receivers, while controlling for caller-receiver relationship type<br>3. Elephants will respond more strongly to playback of call originally addressed to them than to playback of call from same caller originally addressed to another individual |
| 2. Vocal labels are arbitrary (not imitative of receiver's calls) | 1. Calls from caller A to receiver B will be no more similar to receiver B's calls than calls from caller A to other receivers are to receiver B's calls<br>2. Receiver can be predicted from call structure regardless of whether calls are convergent or divergent from receiver's calls relative to other calls by the same caller<br>3. Calls with the same caller and same receiver will be more similar than calls with the same caller and different receivers regardless even among calls that are divergent from the receiver's calls |
| 3. Different callers use same label for same receiver | 1. Receiver ID can be predicted from call structure independently of caller ID<br>2. Calls with different callers and same receiver will be more similar than calls with different callers and different receivers |

**Extended Table 2. Acoustic features used in the random forest models**

| Derived acoustic feature | Derived from | Description |
|---|---|---|
| Median | Sparse matrix | Median of all the values in the sparse matrix |
| Robust skewness | Sparse matrix | Scaled median difference of the left and right half of the distribution of values in the sparse matrix. Estimate of skewness of sparse matrix |
| Minimum extent | Sparse matrix | Estimate of spread of sparse matrix, calculated as sum(x)/max(x)/length(x), where x is the sparse matrix |
| Equivalent statistical extent | Sparse matrix | Estimate of spread of sparse matrix, calculated as sum(x)^2/sum(x^2)/length(x), where x is the sparse matrix |
| Means of low-rank principal components explaining 99.9% of variation | Low-rank matrix | Arithmetic means of the robust principal components required to explain 99.9% of variation in low-rank matrix (74 for spectral features, 12 for cepstral features) |
| Frequencies of unweighted PC spectral peaks 1-4 | Low-rank matrix | Frequencies of the four largest peaks aggregated across the unweighted principal component spectra, sorted from lowest to highest frequency (4 values) |
| Magnitudes of unweighted PC spectral peaks 1-4 | Low-rank matrix | Amplitudes of the four largest peaks aggregated across the unweighted principal component spectra, sorted from lowest to highest frequency (4 values) |
| Frequencies of weighted PC spectral peaks 1-4 | Low-rank matrix | Frequencies of the four largest peaks aggregated across the weighted principal component spectra, sorted from lowest to |

| | | |
|---|---|---|
| | | highest frequency (4 values). Spectra were weighted by the proportion of variation explained by each principal component |
| Magnitudes of weighted PC spectral peaks 1-4 | Low-rank matrix | Amplitudes of the four largest peaks aggregated across the weighted principal component spectra, sorted from lowest to highest frequency (4 values). Spectra were weighted by the proportion of variation explained by each principal component |

All acoustic features were derived from either the sparse matrix or low-rank matrix of a robust principal components analysis performed on multiple acoustic contours of equal length that were measured directly from the signal. For the spectral acoustic features, the acoustic contours were the Hilbert amplitude envelope, the vector of energies in each of the 26 bands of a mel spectrogram, and the delta and delta-delta values of the mel spectral bands. For the cepstral acoustic features, the acoustic contours were the Hilbert amplitude envelope, first 12 mel-frequency cepstral coefficients, and the delta and delta-delta values of the first 12 cepstral coefficients. The principal components analysis was performed on a matrix of all the contours for each call stacked end-to-end.

**Extended Table 3. Definitions of social relationship categories between caller and receiver**

| **Relationship category** | **Code** | **Definition** |
|---|---|---|
| Mother of adult female | Mom2AdDau | Caller is mother of receiver, and receiver is adult female |
| Mother of juvenile | Mom2JuvOff | Caller is mother of receiver, and receiver is juvenile |
| Adult daughter | Adult2Mom | Caller is daughter of receiver and is an adult |
| Juvenile offspring | Juv2Mom | Caller is offspring of receiver and is a juvenile |
| Older adult female to younger adult female | Ad2YngrAd | Caller and receiver are both adult females from same family group, and caller is ≥6 years older than receiver and not mother of receiver |
| Adult (female) age mates | AdAgeMates | Caller and receiver are <6 years apart, from same family group, at least one is an adult female, and neither one is an adult male |
| Younger adult female to older adult female | Ad2OldrAd | Caller is adult female, ≥6 years younger than receiver, not daughter of receiver, and both are from same family group |
| Adult female to juvenile | Adult2Juv | Caller is adult female and not mother of receiver, receiver is juvenile and ≥6 years younger than caller, and both are from same family group |
| Juvenile to adult female | Juv2Adult | Caller is juvenile and ≥6 years younger than caller, receiver is adult female and not mother of caller, and both are from same family group |
| Juvenile to juvenile | Juv2Juv | Caller and receiver are both juveniles from same family group |
| Different groups | FemDiffGrps | Caller and receiver are from different family groups and neither one is adult male |

| | | |
|---|---|---|
| Adult male | AdultMale | Caller and/or receiver is adult male |

Categories were defined based on sex, age, and mother-offspring status, the most important factors influencing dominance and bond strength within an elephant family group. Females were defined as adults if ≥10 years old, and males were defined as adults if independent from their natal group. All non-adults under this definition were classified as juveniles. Six years was chosen as the cutoff for different age classes because it is between 1-2x the average inter-birth interval, so a female ≥6 years older than another individual could have been that individual's allomother.

**Extended Table 4. Results for ANOVAs to test if calls with the same caller and receiver were more similar than calls with the same caller and different receivers**

| Model term | F statistic | DF | *P*-value |
|---|---|---|---|
| ***Spectral acoustic features – all pairs*** | | | |
| Same Caller Pair Type | 94.61 | 1 | <0.0001 |
| Same Context | 23.14 | 1 | <0.0001 |
| ***Spectral acoustic features – convergent calls only*** | | | |
| Same Caller Pair Type | 15.30 | 1 | 0.0001 |
| Same Context | 1.39 | 1 | 0.240 |
| ***Spectral acoustic features – divergent calls only*** | | | |
| Same Caller Pair Type | 8.67 | 1 | 0.0033 |
| Same Context | 13.22 | 1 | 0.0003 |
| ***Cepstral acoustic features – all pairs*** | | | |
| Same Caller Pair Type | 42.09 | 1 | <0.0001 |
| Same Context | 34.71 | 1 | <0.0001 |
| ***Cepstral acoustic features – convergent calls only*** | | | |
| Same Caller Pair Type | 26.40 | 1 | <0.0001 |
| Same Context | 0.005 | 1 | 0.946 |
| ***Cepstral acoustic features – divergent calls only*** | | | |
| Same Caller Pair Type | 5.37 | 1 | 0.021 |
| Same Context | 33.03 | 1 | <0.0001 |

Each observation was a pair of calls. ANOVA models were of the form Rank-transformed Proximity Score ~ Same Caller Pair Type (whether the two calls in a pair had the same caller and receiver or same caller and different receivers) + Same Context (whether the two calls in a pair had the same behavioral context). Pairs of calls recorded on the same date or where the two calls had a different type of caller-receiver relationship were excluded. Three models were run for each set of acoustic features (spectral and cepstral): all pairs of calls meeting above criteria (n=2391), pairs of calls in which both calls were convergent on the receiver's calls (n=252), and pairs of calls in which both calls were divergent from the receiver's calls (n=798). Convergent calls = calls from caller A to receiver B that were more similar to receiver B's calls than calls from caller A to other receivers were to receiver B's calls. Divergent calls = calls from caller A to receiver B that were less similar to receiver B's calls than calls from caller A to other receivers were to receiver B's calls.

**Extended Table 5. Results for ANOVAs to test if calls addressed to a given receiver were imitative of the receiver's calls**

| Model term | Cohen's D | F statistic | DF | *P*-value |
|---|---|---|---|---|
| ***Spectral acoustic features*** | | | | |
| Imitation Pair Type | 0.0037 | 11.70 | 1 | 0.0006 |
| Same Relationship | | 0.66 | 1 | 0.416 |
| Same Context | | 5.27 | 1 | 0.022 |
| Caller Dyad ID | | 13.16 | 112 | <0.0001 |
| ***Cepstral acoustic features*** | | | | |
| Imitation Pair Type | 0.049 | 24.58 | 1 | <0.0001 |
| Same Relationship | | 1.89 | 1 | 0.169 |
| Same Context | | 14.02 | 1 | 0.0002 |
| Caller Dyad ID | | 8.24 | 112 | <0.0001 |

Each observation was a pair of calls. ANOVA models were of the form Rank-transformed Proximity Score ~ Imitation Pair Type + Same Relationship + Same Context + Caller Dyad ID. Model was run once for each set of acoustic features: spectral and cepstral. Imitation Pair Type = whether the caller of one call in a pair was the receiver of the other call. Same Relationship = whether the callers of both calls in a pair had the same type of relationship to their respective receivers. Same Context = whether the two calls in a pair were recorded in the same behavioral context (contact/greeting). Caller Dyad ID = identifier for the two callers in a pair. Pairs of calls recorded on the same date, from callers in different social groups, or with the same caller or receiver were excluded. We also excluded pairs of calls for which Caller Dyad ID only occurred with one level of Imitation Pair Type (final n=11,309).

**Extended Table 6. Results for ANOVAs to test if different callers used similar labels for the same receiver**

| **Model term** | **F statistic** | **DF** | ***P*-value** |
|---|---|---|---|
| ***Spectral acoustic features*** | | | |
| Different Caller Pair Type | 74.96 | 1 | <0.0001 |
| Same Relationship | 42.41 | 1 | <0.0001 |
| Same Context | 24.19 | 1 | <0.0001 |
| ***Cepstral acoustic features*** | | | |
| Different Caller Pair Type | 5.68 | 1 | 0.017 |
| Same Relationship | 89.35 | 1 | <0.0001 |
| Same Context | 68.45 | 1 | <0.0001 |

Each observation was a pair of calls. ANOVAs were of the form Rank-transformed Proximity Score ~ Different Caller Pair Type + Same Relationship + Same Context. Model was run separately for each set of acoustic features: spectral and cepstral. Different Caller Pair Type = whether the two calls in a pair had different callers and the same receiver or different callers and different receivers. Same Relationship = whether the two calls in a pair had the same type of relationship between caller and receiver. Same Context = whether the two calls in a pair were recorded in the same behavioral context (contact/greeting) or not. Pairs of calls recorded on the same date or from callers in different social groups were excluded (final n=20,235).

**Extended Table 7. Summary of playback trials for each subject**

| Subject | Date | Treatment | Original caller | Original receiver | Caller's relationship to orig. recvr | Number previous attempts |
|---|---|---|---|---|---|---|
| R24 | 2021-10-18 | Control | R23 (a) | R23.09 | Mother | 1 |
| R24 | 2021-11-18 | Test | R23 (b) | R24 | Age mates | 1 |
| R23.09 | 2021-11-27 | Control | R23 (b) | R24 | Age mates | 1 |
| R23.09 | 2022-03-31 | Test | R23 (a) | R23.09 | Mother | 2 |
| R22.89 | 2021-10-21 | Control | R24 (a) | R23.93 | Older | 0 |
| R22.89 | 2021-10-28 | Test | R24 (b) | R22.89 | Older | 0 |
| R23.93 | 2021-10-25 | Test | R24 (a) | R23.93 | Older | 0 |
| R23.93 | 2022-04-05 | Control | R24 (b) | R22.89 | Older | 2 |
| R23 | 2022-02-03 | Test | R24 (c) | R23 | Age mates | 0 |
| R23 | 2022-02-22 | Control | R24 (d) | R21.04 | Older | 0 |
| R21.04 | 2022-03-26 | Control | R24 (c) | R23 | Age mates | 2 |
| R21.04 | 2022-04-21 | Test | R24 (d) | R21.04 | Older | 1 |
| R26 | 2021-10-23 | Control | R25.03 (a) | R25.9002 | Age mates | 0 |
| R26 | 2021-11-20 | Test | R25.03 (b) | R26 | Younger | 0 |
| R25.9002 | 2021-11-02 | Test | R25.03 (a) | R25.9002 | Age mates | 0 |
| R26.00 | 2021-12-11 | Test | R25.03 (c) | R26.00 | Age mates | 0 |
| R26.00 | 2021-12-18 | Control | R25.03 (b) | R26 | Younger | 0 |
| R26.00 | 2022-03-16 | Control | R26 (a) | R25.03 | Older | 0 |
| R26.00 | 2022-04-06 | Test | R26 (b) | R26.00 | Mother | 1 |
| R25.03 | 2022-03-25 | Control | R26 (b) | R26.00 | Mother | 0 |
| R25.03 | 2022-04-04 | Test | R26 (a) | R25.03 | Older | 2 |
| M25.0012 | 2021-11-05 | Test | M25.00 (a) | M25.0012 | Mother | 0 |
| M25.0012 | 2022-03-24 | Control | M25.00 (b) | M25.0017 | Mother | 2 |
| M6.99 | 2022-03-01 | Test | M6 (a) | M6.99 | Mother | 0 |
| M6.99 | 2022-03-21 | Control | M6 (b) | M7.8905 | Older | 0 |
| M7.8905 | 2022-03-01 | Control | M6 (a) | M6.99 | Mother | 0 |
| M7.8905 | 2022-03-08 | Test | M6 (b) | M7.8905 | Older | 0 |
| M8.88 | 2022-03-15 | Control | M8.08 (a) | M8.98 | Younger | 1 |
| M8.88 | 2022-03-22 | Test | M8.08 (b) | M8.88 | Younger | 1 |
| M8.98 | 2022-01-29 | Test | M8.08 (a) | M8.98 | Younger | 0 |
| M8.98 | 2022-02-08 | Control | M8.08 (b) | M8.88 | Younger | 0 |
| M9 | 2021-12-18 | Test | M9.96 (a) | M9 | Daughter | 0 |
| M9 | 2022-04-13 | Control | M9.96 (b) | M9.9612 | Mother | 0 |
| M9.9612 | 2022-02-21 | Control | M9.96 (a) | M9 | Daughter | 0 |

All callers and subjects were adult females except M25.0012 (subadult male) and M9.9612 (subadult female). The letter in parentheses after each caller ID represents a unique call (e.g., R23 (a) and R23 (b) were different calls recorded from R23). Twelve trials were redone once or twice because the playback system malfunctioned, the subject went out of sight just as the playback began, or the speaker was accidentally placed >60 m away or in the wrong direction relative to the subject and the original caller. Trials that were later redone are not included in this table.
